# Uterine progesterone signaling is a target for metformin therapy in PCOS-like rats

**DOI:** 10.1101/238261

**Authors:** Min Hu, Yuehui Zhang, Jiaxing Feng, Xue Xu, Jiao Zhang, Wei Zhao, Xiaozhu Guo, Juan Li, Edvin Vestin, Peng Cui, Xin Li, Xiao-ke Wu, Mats Brännström, Linus R Shao, Håkan Billig

## Abstract

Impaired progesterone (P4) signaling is linked to endometrial dysfunction and infertility in women with polycystic ovary syndrome (PCOS). Here we report for the first time that elevated expression of progesterone receptor (PGR) isoforms A and B parallels increased estrogen receptor (ER) expression in PCOS-like rat uteri. The aberrant PGR-targeted gene expression in PCOS-like rats before and after implantation overlaps with dysregulated expression of *Fkbp52* and *Ncoa2*, two genes that contribute to the development of uterine P4 resistance. *In vivo* and *in vitro* studies of the effects of metformin on the regulation of the uterine P4 signaling pathway under PCOS conditions showed that metformin directly inhibits the expression of PGR and ER along with the regulation of several genes that are targeted dependently or independently of PGR-mediated uterine implantation. Functionally, metformin treatment corrected the abnormal expression of cell-specific PGR and ER and some PGR-target genes in PCOS-like rats with implantation. Additionally, we documented how metformin contributes to the regulation of the PGR-associated MAPK/ERK/p38 signaling pathway in the PCOS-like rat uterus. Our data provide novel insights into how metformin therapy regulates uterine P4 signaling molecules under PCOS conditions.

## Introduction

Polycystic ovary syndrome (PCOS) is a clinically and etiologically heterogeneous hormone-imbalance disorder that is associated with multiple reproductive and metabolic abnormalities (Rosenfield and Ehrmann 2016). Women suffering from PCOS present with arrested folliculogenesis and chronic anovulation-linked infertility (Azziz *et al.* 2016; Rosenfield and Ehrmann 2016), and they also have more adverse reproductive risk as evidenced by an increase in the prevalence of implantation failure, recurrent miscarriage, spontaneous abortion, premature delivery, endometrial carcinoma (Goodarzi *et al.* 2011; Palomba *et al.* 2015; Shao *et al.* 2014b). In addition to the ovarian dysfunction (Azziz *et al.* 2016; Rosenfield and Ehrmann 2016), it is assumed that the impairment of endometrial function also contributes to PCOS-associated infertility (Evans *et al.* 2016). Although progesterone (P4)-based oral contraceptive therapy is often efficacious (Lopes *et al.* 2014; Vrbikova and Cibula 2005), perturbations in endometrial P4 signaling that result from attenuated responsiveness and resistance to P4 are common in the endometrium of a PCOS patient (Li *et al.* 2014a; Piltonen 2016). P4 resistance is a condition in which tissues and cells do not respond appropriately to P4 (Chrousos *et al.* 1986), and this is evidenced by endometriosis and endometrial hyperplasia that may progress to endometrial carcinoma despite supplementation with P4 or its analogs (Gunderson *et al.* 2012; Shao *et al.* 2014a). Gene profiling experiments have shown that different endometrial genes are likely to act in concert in this abnormal condition in PCOS patients (Kim *et al.* 2009; Savaris *et al.* 2011); however, how changes in the expression of P4 signaling molecules contribute to the P4 resistance in a PCOS patient’s endometrium is poorly understood.

P4 is an essential contributing factor in female reproductive tissues that regulates multiple physiological processes such as the menstrual cycle, implantation, pregnancy maintenance, and labor initiation (Evans *et al.* 2016). There are two major progesterone receptor (PGR) isoforms, PGRA and PGRB, both of which are involved in a common P4 signaling pathway for uterine cell-specific proliferation and differentiation (Li *et al.* 2014a; Patel *et al.* 2015). P4 binding activates both PGR isoforms and leads to translocation from the cytosol to the nucleus followed by binding to the P4-responsive elements of the target genes, resulting in alterations of PGR-targeted gene expression depending on the recruitment of co-regulators (Patel *et al.* 2015). It has been reported that endometrial PGR expression is elevated in PCOS patients who have anovulation compared to PCOS patients who still ovulate and to non-PCOS patients (Margarit *et al.* 2010; Quezada *et al.* 2006). Additionally, PGR activity is also modulated by the cytoplasmic mitogen-activated protein kinase (MAPK)/extracellular signal-regulated kinase (ERK) signaling pathway (Gellersen and Brosens 2014; Patel *et al.* 2015). While high levels of ERK1/2 expression and activation reflect the P4-PGR signaling-induced decidualization status in human and rodent uteri (Lee *et al.* 2013; Tapia-Pizarro *et al.* 2017; Thienel *et al.* 2002), it remains to be determined whether suppression of MAPK/ERK signaling occurs in the endometrium and whether such dysregulation can negatively impact uterine function under PCOS conditions.

Metformin is an anti-diabetic drug that is a clinically approved treatment in PCOS patients worldwide (Naderpoor *et al.* 2015). Several diverse molecular mechanisms of metformin have been demonstrated in human endometrial carcinoma tissues *in vivo* and in different endometrial cancer cells *in vitro* (Shao *et al.* 2014b), and metformin’s therapeutic effects on endometrial function are evidenced by improvement of endometrial receptivity, enhancement of endometrial vascularity and blood flow, and reversion of endometrial hyperplasia and carcinoma into normal endometria in some women with PCOS (Jakubowicz *et al.* 2001; Li *et al.* 2014b; Palomba *et al.* 2006). Our recent studies using a PCOS-like rat model found that chronic treatment with metformin has significant anti-androgenic and anti-inflammatory impacts in the uterus (Zhang *et al.* 2017). Given the central role of P4 signaling in uterine implantation (Evans *et al.* 2016; Patel *et al.* 2015) and the ability of metformin to rescue implantation failure in some PCOS-like rats by modulating the expression of multiple implantation-related genes in the uterus *in vivo* (Zhang *et al.* 2017), we speculated that the beneficial effects of metformin might be mechanistically linked to the uterine P4 signaling pathway under pathological conditions such as PCOS. To address this hypothesis, we analyzed PCOS-associated PGR isoform expression and the MAPK signaling network in human and rat uterine tissues. By combining a PCOS-like rat model (Zhang *et al.* 2016) and *in vitro* tissue culture approach (Li *et al.* 2015), we aimed to determine whether metformin directly reverses aberrant PGR-targeted and implantation-related gene expression in the PCOS-like rat uterus.

## Materials and Methods

### Study approval

All animal experiments were performed according to the National Institutes of Health guidelines on the care and use of animals and were approved and authorized by the Animal Care and Use Committee of the Heilongjiang University of Chinese Medicine, China (HUCM 2015-0112).

### Experimental animals and tissue preparations

Adult female Sprague–Dawley rats (n = 134) were obtained from the Laboratory Animal Centre of Harbin Medical University, Harbin, China (License number SCXK 2013-001). Animals were housed in the animal care facility with free access to food and water and a controlled temperature of 22°C ± 2°C with a 12 h light/dark cycle. Estrous cycles were monitored daily by vaginal lavage according to a standard protocol (Feng *et al.* 2010). All rats (70 days old) with the different stages of estrous cycle used in this study were confirmed by examination of vaginal smears under a light microscope for two sequential cycles (about 8–10 days). Any PCOS-like (insulin+hCG-treated) rats that exhibited prolonged estrous cycles (more than 5 days) were excluded from the study.

Experiment 1: Rats were randomly divided into control (saline treatment, n = 20) and experimental (PCOS-like, n = 20) groups. The experimental group was treated with insulin plus hCG to induce a PCOS-like metabolic and reproductive phenotype, and the control rats were treated with an equal volume of saline (Zhang *et al.* 2018; Zhang *et al.* 2016). In brief, insulin was started at 0.5 IU/day and gradually increased to 6.0 IU/day between day 1 and the day 22 to induce hyperinsulinemia and insulin resistance, and 3.0 IU/day hCG was given on all 22 days to induce hyperandrogenism. Animals were treated with twice-daily subcutaneous injections until the end of the experiment. Rats with repeated insulin injections have not shown any hypoglycemic episodes (Bogovich *et al.* 1999; Damario *et al.* 2000; Poretsky *et al.* 1992; Zhang *et al.* 2018). Detailed analysis of endocrine and metabolic parameters as well as the uterine morphology in these animals has been reported previously (Zhang *et al.* 2016). On day 23, each group of rats was divided into two subgroups of 10 rats each (Supplemental Fig. 1A). For treatment, metformin was dissolved in saline and given as a daily oral dose of 500 mg/kg by a cannula. The treatment time and tissue collection are described in our previous study (Zhang *et al.* 2017).

Experiment 2: Rats were randomly divided into control (saline treatment, n = 21) and experimental (PCOS-like, n = 15) groups and treated as described in Experiment 1 (Supplemental Fig. 1B). After metformin treatment, control and PCOS-like rats were mated with fertile males of the same strain to induce implantation, which was determined by the presence of a vaginal plug (day 1 of pregnancy). The rats were sacrificed between 0800 and 0900 hours on day 6 of pregnancy. To identify the implantation sites, rats were injected intravenously with a Chicago Blue B dye solution (1% in saline) and sacrificed 10 min later. Uteri were dissected and assessed for clearly delineated blue bands as evidence of early implantation sites as described previously (Zhang *et al.* 2017).

Experiment 3: Rats were divided into control (saline treatment, n = 9) and experimental (PCOS-like, n = 39) groups and treated as described in the Experiment 1. On the 23rd day, the PCOS-like rats were divided into four subgroups and treated daily with P4 (4 mg/kg), RU486 (6 mg/kg), or both for 3 days. For treatment, P4 and RU486 were dissolved in 100% ethanol and resuspended in sesame oil. All subcutaneous injections were in a volume of 100 µl. An equal volume of 100% ethanol and sesame oil was injected into both healthy control rats and PCOS-like rats as experimental controls (Supplemental Fig. 1C). The pharmacological doses and treatment time intervals of P4 and RU486 were chosen on the basis of previous studies (Kim *et al.* 2006; Knox *et al.* 1996).

After dissection, the uterine horns were trimmed free of fat and connective tissue. One side of the uterus in each animal was fixed in 10% neutral formalin solution for 24 h at 4°C and embedded in paraffin for histochemical analysis. The other side was immediately frozen in liquid nitrogen and stored at -70°C for Western blot and quantitative real-time PCR (qRT-PCR) analysis.

Detailed description of the methods including the primary *in vitro* tissue culture and treatment, morphological assessment and immunostaining, protein isolation and Western blot analysis, RNA extraction and qRT-PCR analysis, and measurement of biochemical parameters used in this study are provided in Supplemental files.

### Statistical analysis

GraphPad Prism was used for statistical analysis and graphing. For all experiments, n-values represent the number of individual animals. Data are represented as the means ± SEM. Statistical analyses were performed using SPSS version 24.0 statistical software for Windows (SPSS Inc., Chicago, IL). The normal distribution of the data was tested with the Shapiro–Wilk test. Differences between groups were analyzed by one-way ANOVA or two-way ANOVA, and this was followed by Tukey’s post-hoc test for normally distributed data or the Kruskal–Wallis test followed by the Mann–Whitney U-test for skewed data. All *p*-values less than 0.05 were considered statistically significant.

## Results

### Metformin alters PGR isoform and PGR-targeted gene expression in PCOS-like rats

The insulin+hCG-treated rats exhibit reproductive disturbances that mimic human PCOS (Zhang *et al.* 2017; Zhang *et al.* 2016). Prompted by these findings, we set out to investigate the impact of P4 signaling in this model. First, we showed that although the ratio of PGRA to PGRB was not significantly different between control and PCOS-like rats, the PCOS-like rats had increased levels of uterine PGRA and PGRB (Fig. 1A). While PGR immunoreactivity was primarily evidenced in control rat uterine luminal and glandular epithelia as well as in the stroma, the immunoreactivity of luminal epithelial PGR expression was associated with increased numbers of luminal epithelial cells and increased immunoreactivity of PGR in the stroma in PCOS-like rats (Fig. 1B). Metformin treatment did not significantly affect PGR isoform expression in control rats and PCOS-like rats compared to those rats treated with saline (Fig. 1A). However, we found that PGR immunoreactivity was decreased in the luminal and glandular epithelia by metformin treatment in both control rats and PCOS-like rats compared to those treated with saline (Fig. 1B). Conversely, intense immunoreactivity of PGR expression was detected in the stroma located close to the luminal epithelia in control and PCOS-like rats treated with metformin (Fig. 1B). In contrast to the epithelia and stroma, no significant changes in PGR expression in the myometrium were found in any of the groups (data not shown). Because a large body of evidence indicates that regulation of P4 signaling results in changes in the expression of several PGR-targeted genes in the uterus (Bhurke *et al.* 2016), we profiled the expression of genes that are indicators for PGR activity in the rat uterus by qRT-PCR. Quantitative data indicated that *Smo*, and *Nr2f2* mRNA levels were increased in PCOS-like rats compared to control rats treated with saline. In contrast, the *Fkbp52* mRNA level was decreased in PCOS-like rats compared to control rats (Fig. 1C). We next determined the actions of metformin treatment on PGR-targeted gene expression and showed that *Ptch, Fkbp52*, and *Ncoa2* levels were increased in PCOS-like rats treated with metformin compared to PCOS-like rats treated with saline, while *Smo* and *Nr2f2* mRNA levels were decreased on PCOS-like rats treated with metformin compared to those treated with saline (Fig. 1C).

**Figure 1.**
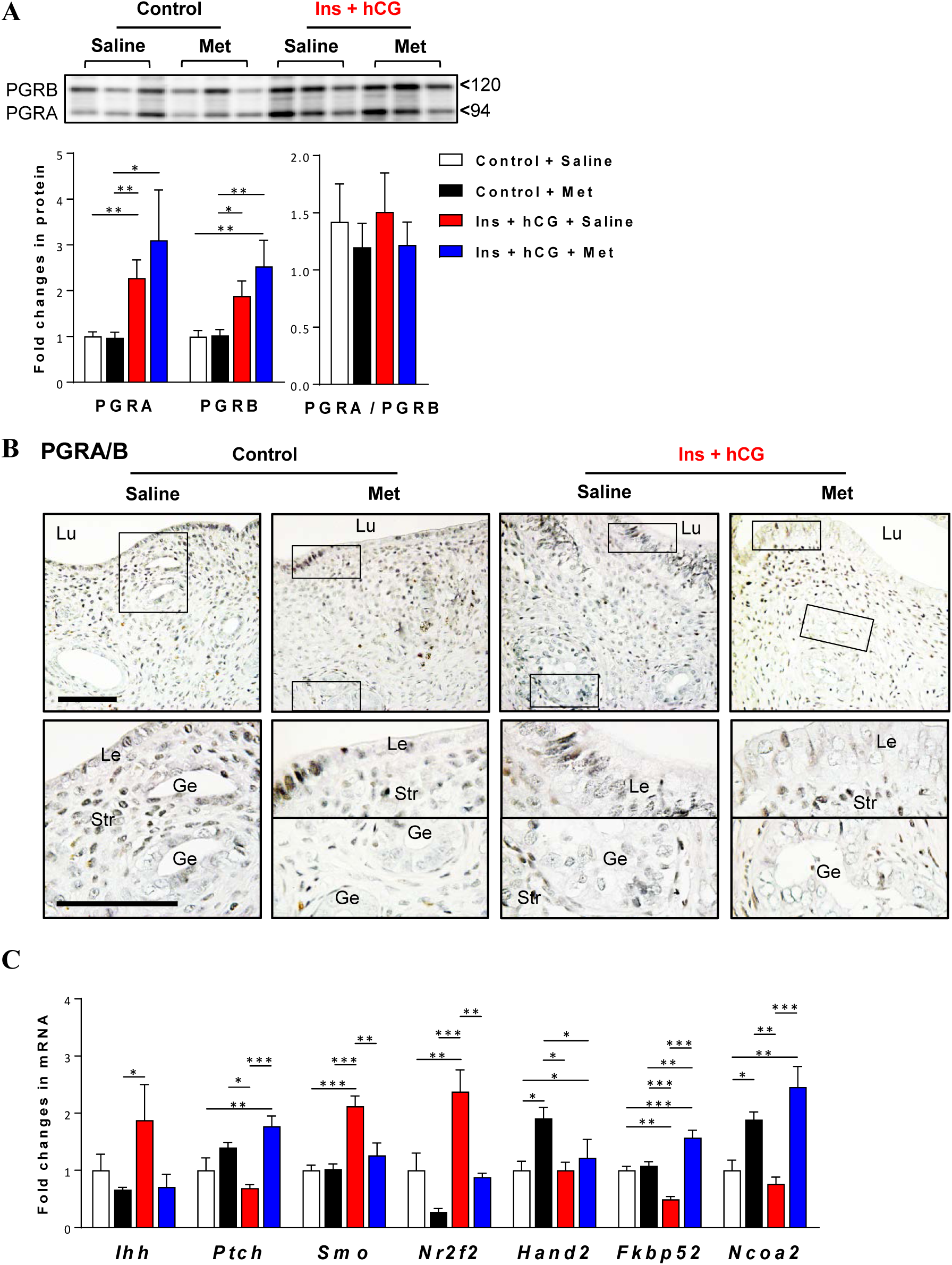
Chronic treatment with metformin alters PGR isoform protein expression and PGR-target gene expression in the rat uterus *in vivo*. A, Western blot analysis of protein expression in the rat uterus was performed. Representative images and quantification of the densitometric data (n = 8–9/group) of PGR isoforms are shown. B, Immunohistochemical detection of PGRA/B in control rats treated with saline or metformin and in insulin+hCG-treated rats treated with saline or metformin. Representative images (n = 5/group) are shown. Lu, lumen; Le, luminal epithelial cells; Ge, glandular epithelial cells; Str, stromal cells. Scale bars (100 µm) are indicated in the photomicrographs. High magnification images are shown in the bottom panels. C, Uterine tissues from control rats treated with vehicle or metformin and insulin+hCG-treated rats treated with saline or metformin (n = 6/group) were analyzed for mRNA levels of *Ihh, Ptch, Smo, Nr2f2, Hand2, Fkbp52*, and *Ncoa2* by qRT-PCR. The mRNA level of each gene relative to the mean of the sum of the *Gapdh* and *U87* mRNA levels in the same sample is shown. Values are expressed as means ± SEM. Statistical tests are described in the Material and Methods. * *p* < 0.05; ** *p* < 0.01; *** *p* < 0.001.

### Metformin partially prevents implantation failure in parallel with regulation of PGR isoform and PGR-targeted gene expression in PCOS-like rats

Metformin has been shown to partially rescue the disruption of the implantation process in PCOS-like rats (Zhang *et al.* 2017), and the altered endocrine and metabolic parameters in these animals are shown in Supplemental Table 4. After metformin treatment, total testosterone levels, the ratio of total testosterone to androstenedione, and fasting insulin levels were all significantly higher in PCOS-like rats where implantation did not occur compared to those with implantation, as was insulin resistance as assessed by the homeostasis model assessment of insulin resistance, mirroring the endocrine and metabolic abnormalities in PCOS patients (Azziz *et al.* 2016; Rosenfield and Ehrmann 2016). Of note, PCOS-like rats that failed to implant embryos also exhibited decreased P4 levels. These data suggest that implantation failure in PCOS-like rats treated with metformin is due not only to hyperandrogenism and insulin resistance, but also to impairment of P4 signaling in the uterus. Further morphological characterization of metformin-treated PCOS-like rats with no implantation revealed the infiltration of immune cells into the glandular epithelial cell layer in a similar manner to when hormone imbalances were studied in a previous report (Wira *et al.* 2005) (Supplemental Fig. 3, black arrowheads). To determine how impairment of P4 signaling causes implantation failure, we subsequently analyzed PGR isoform and PGR-targeted gene expression in PCOS-like rats with no implantation. Although treatment with metformin increased PGR isoform expression in control and PCOS-like rats, neither the PGRA nor PGRB protein level was altered between PCOS-like rats with implantation and with failed implantation (Fig. 2A). As shown in Figure 2B, while PGR protein was expressed in the decidualizing stroma at the site of implantation in all groups, PGR immunoreactivity was increased in the stroma of the inter-implantation region in control rats treated with metformin. Furthermore, we found that the immunoreactivity of PGR was increased in the epithelia in PCOS-like rats without implantation despite metformin treatment. Thus, metformin appeared to participate in the regulation of uterine PGR expression in a cell type-specific manner in PCOS-like rats before and after implantation. qRT-PCR data indicated that *Ihh* and *Ncoa2* mRNAs were increased and that *Ptch* and *Fkbp52* mRNAs were decreased in metformin-treated PCOS-like rats with no implantation compared to control rats treated with saline or metformin and to metformin-treated PCOS-like rats with implantation (Fig. 2C).

**Figure 2.**
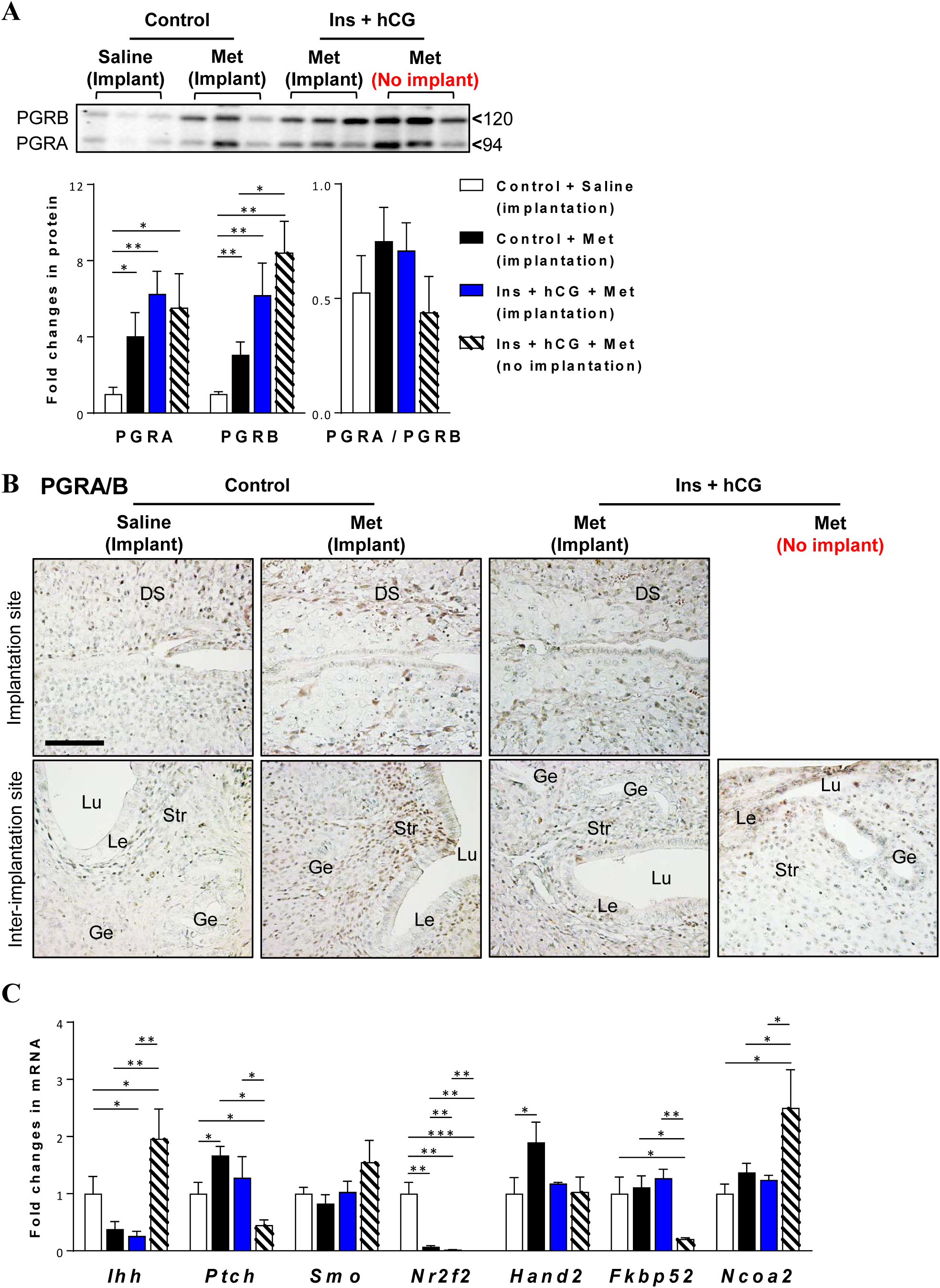
Chronic treatment with metformin alters PGR isoform protein expression and PGR-target gene expression in the rat uterus after implantation. A, Western blot analysis of protein expression in the rat uterus was performed. Representative images and quantification of the densitometric data of PR isoforms are shown (n = 5–7/group). B, Immunohistochemical detection of PGRA/B in the uterine implantation and inter-implantation sites. Representative images are shown (n = 5/group). DS, decidualized stroma; Lu, lumen; Le, luminal epithelial cells; Ge, glandular epithelial cells; Str, stromal cells. Scale bars (100µm) are indicated in the photomicrographs. C, Uterine tissues (n = 5–6/group) were analyzed for mRNA levels of *Ihh, Ptch, Smo, Nr2f2, Hand2, Fkbp52*, and *Ncoa2* by qRT-PCR. The mRNA level of each gene is shown relative to the mean of the sum of the *Gapdh* and *U87* mRNA levels in the same sample. Values are expressed as means ± SEM. Statistical tests are described in the Materials and Methods. * *p* < 0.05; ** *p* < 0.01; *** *p* < 0.001.

**Figure 3.**
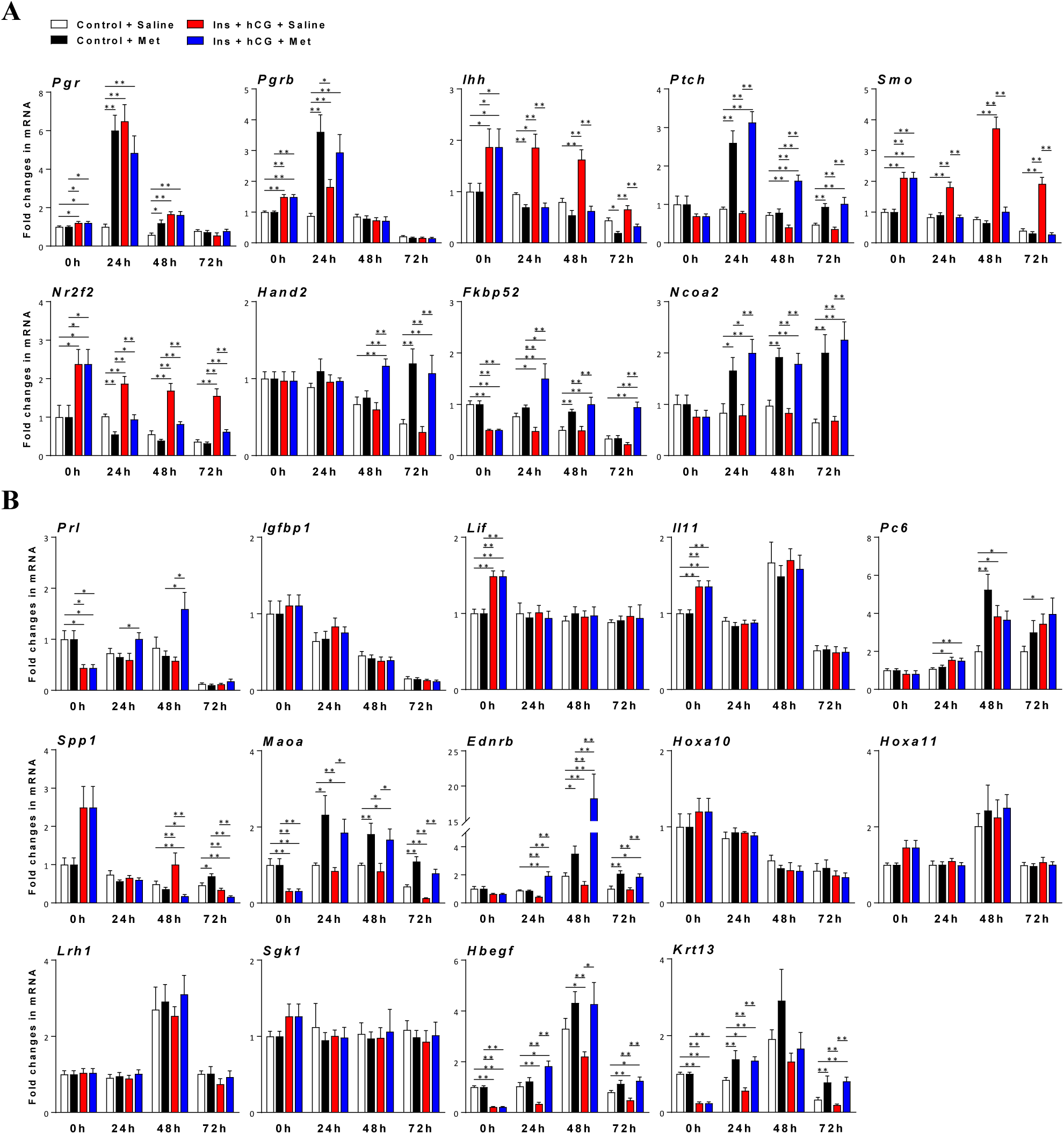
Specific regulation of uterine PGR isoforms, PGR-targeted and implantation-related gene expression by metformin treatment *in vitro*. Quantitative RT-PCR analysis of (A) *Pgr, Pgrb, Ihh, Ptch, Smo, Nr2f2, Hand2, Fkbp52*, and *Ncoa2* and (B) *Prl, Igfbp1, Lif, Il11, Pc6, Spp1, Maoa, Ednrb, Hoxa10, Hoxa11, Lrh1, Sgk1, Hbegf*, and *Krt13* mRNA levels in rat uterine tissues treated with either saline or 10 mM metformin for the indicated culture times (n = 6/group). mRNA levels were normalized to the average levels of *Gapdh* and *U87* mRNA in the same sample. Values are expressed as means ± SEM. Statistical tests are described in the Materials and Methods. * *p* < 0.05; ** *p* < 0.01; *** *p* < 0.001.

### Metformin directly regulates PGR isoform, PGR-target, and implantation-related gene expression in vitro

Based on these *in vivo* observations, we asked whether the effect of metformin was direct or indirect in the PCOS-like rat uterus. *In vitro* uterine tissue culture experiments revealed that *Pgr* and *Pgrb*, mRNA levels were higher in PCOS-like rats compared to control rats, in agreement with alteration of PGR isoform protein expression (Fig. 1C). Furthermore, metformin treatment increased *Pgr* and *Pgrb* mRNA levels in control and PCOS-like rats in a time-dependent manner (Fig. 3A). Consistent with the *in vivo* effects of metformin in PCOS-like rats (Fig. 1C), *Ihh, Smo*, and *Nr2f2* mRNA levels were increased in the PCOS-like rat uterus compared to the control rat uterus and were down-regulated by metformin treatment *in vitro*. While the *Hand2* mRNA level was upregulated by metformin treatment at 48 h and 72 h,we detected the upregulation of *Ptch, Fkbp52*, and *Ncoa2* mRNA levels in the PCOS-like rat uterine tissues over a 72-h course after metformin treatment (Fig. 3A).

The expression of a number of implantation-related genes has been reported to be regulated by metformin treatment in PCOS-like rats during implantation (Zhang *et al.* 2017). These previous observations prompted further analysis of implantation-related gene expression by metformin treatment *in vitro*. In contrast to the different regulation patterns of *Spp1, Lrh1, Sgk1*, and *Krt13* mRNAs under *in vivo* and *in vitro* conditions, the *in vitro* responses of uterine *Prl, Igfbp1, Il11, Pc6, Maoa, Ednrb, Hoxa10, Hoxa11*, and *Hbegf* mRNA levels to metformin (Fig. 3B) were coincident with the *in vivo* regulation of the expression pattern of these genes (Zhang *et al.* 2017). Our data indicated that metformin directly up-regulates uterine *Prl, Maoa, Ednrb*, and *Hbegf* mRNA levels in PCOS-like rats during implantation *in vivo*.

To ascertain whether the modulation of uterine gene expression is P4-mediated and PGR-dependent in PCOS-like rats, insulin+hCG-treated rats were injected subcutaneously with P4 and/or RU 486 for three days. As shown in Figure 4, the increased PGR isoform protein levels (Fig. 1A) were confirmed by analysis of *Pgr* and *Pgrb* mRNA expression in the PCOS-like rat uterus. Although treatment with P4 and/or RU486 did not significantly affect *Pgr* mRNA expression, we found that *Pgrb* mRNA levels were decreased in PCOS-like rats compared to those rats with no treatment (Fig. 4). Among seven PGR-targeted genes (Fig. 1C), we found that *Ptch, Hand2*, and *Fkbp52* mRNA levels were increased and that *Ihh, Smo*, and *Nr2f2* mRNA levels were decreased in PCOS-like rats treated with P4 compared to those rats with no treatment. We also observed that treatment with RU486 alone or combined with P4 reversed the changes in *Smo, Hand2*, and *Fkbp52* mRNA levels in PCOS-like rats (Fig. 4). No significant differences of uterine *Ncoa2* mRNA expression were observed in PCOS-like rats regarding the different treatments. Based on our current experimental approaches, it is likely that another regulatory mechanism contributes to the metformin-induced up-regulation of *Ncoa2* mRNA levels in PCOS-like rats.

**Figure 4.**
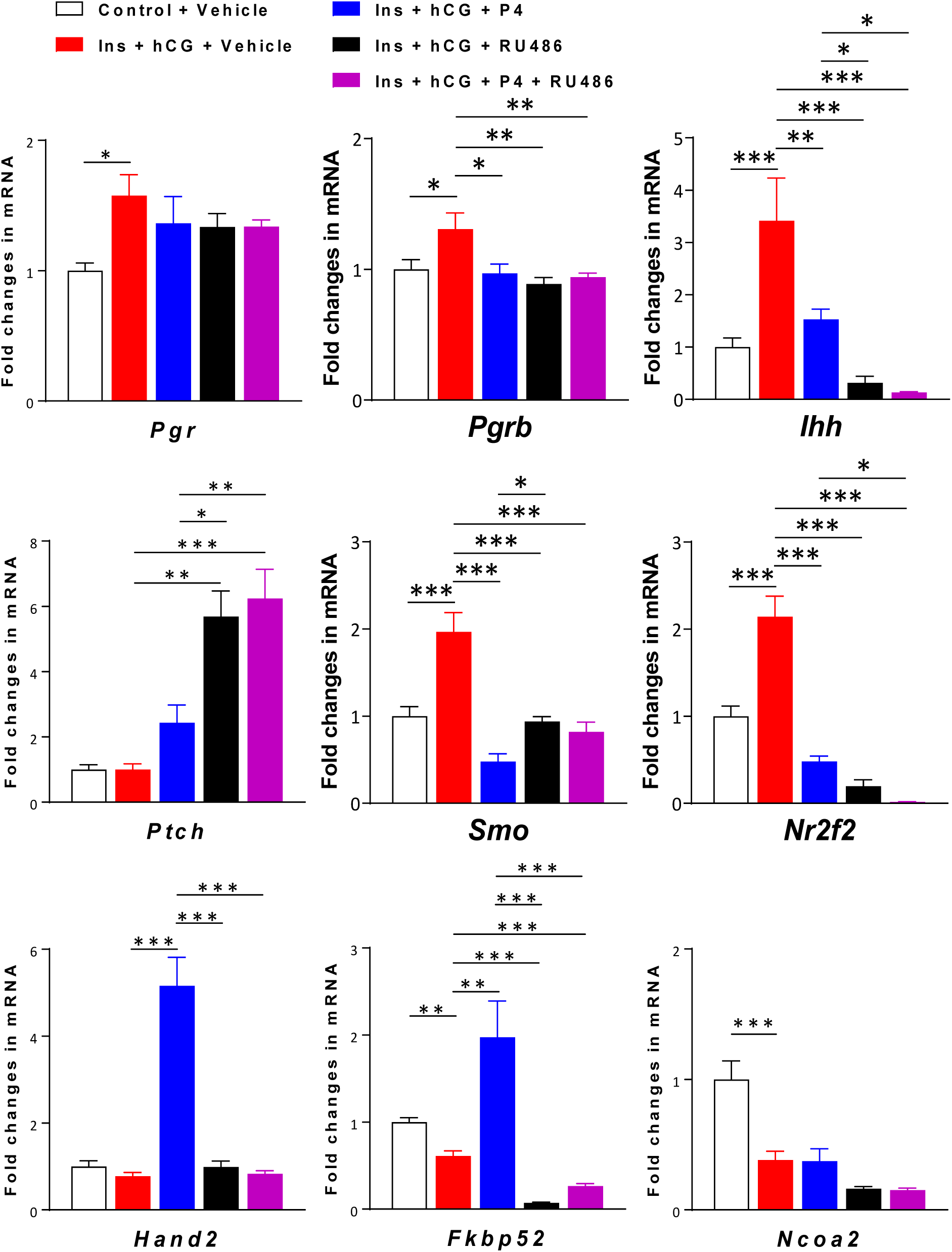
Specific regulation of uterine PGR isoforms and PGR-targeted gene expression by treatment with P4 and/or RU486 *in vivo*. Uterine tissues (n = 5–6/group) were analyzed for mRNA levels of *Pgr, Pgrb, Ihh, Ptch, Smo, Nr2f2, Hand2, Fkbp52*, and *Ncoa2* by qRT-PCR. mRNA levels were normalized to the average levels of *Gapdh* and *U87* mRNA in the same sample. Values are expressed as means ± SEM. Statistical tests are described in the Materials and Methods. * *p* < 0.05; ** *p* < 0.01; *** *p* < 0.001.

### Metformin regulates the MAPK signaling pathway in PCOS-like rats before and after implantation

In an attempt to understand the changes in PGR activation and function observed in PCOS patients (Patel *et al.* 2015), we performed a Western blot analysis to measure the expression of several proteins that are involved in the MAPK signaling pathway in the uterus after metformin treatment. As shown in Figure 5A, there was no significant difference in p-c-Raf, p-MEK1/2, p-ERK1/2, p-p38 MAPK, or p38 MAPK expression between saline-treated and metformin-treated rats. Quantitative protein data indicated that the expression of p-p38 MAPK and p38 MAPK was significantly decreased in PCOS-like rats compared to control rats. Nevertheless, metformin treatment only reversed p-p38 MAPK protein expression in PCOS-like rats.

**Figure 5.**
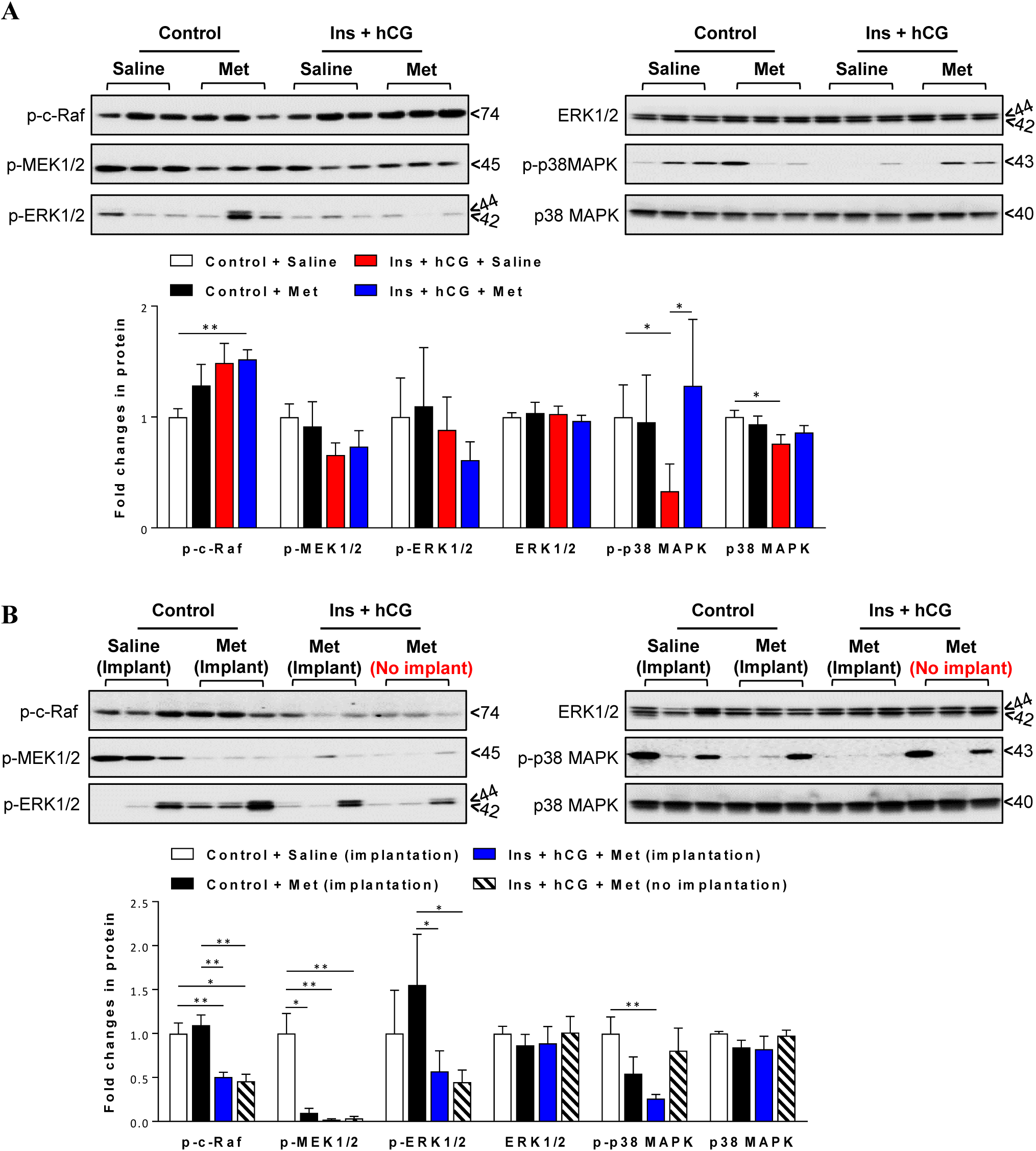
Chronic treatment with metformin alters the MAPK signaling pathway in the rat uterus before and after implantation. Western blot analysis of protein expression in the rat uterus was performed. Representative images and quantification of the densitometric data for p-c-Raf, p-MEK1/2, p-ERK1/2, ERK1/2, p-p38 MAPK, and p38 MAPK are shown (n = 6–9/group before implantation; n = 5–6/group after implantation). Values are expressed as means ± SEM. Statistical tests are described in the Materials and Methods. * *p* < 0.05; ** *p* < 0.01.

We next assessed whether the MAPK/ERK signaling pathway contributes to uterine implantation in control and PCOS-like rats treated with metformin. As shown in Figure 5B, although the p-MEK1/2 level was decreased in control rats treated with metformin compared to control rats treated with saline, no significant difference in p-c-Raf, p-ERK1/2, ERK1/2, p-p38 MAPK, or p38 MAPK expression between these two groups was found. Furthermore, our data showed that p-c-Raf, p-MEK1/2, and p-ERK1/2 protein levels were down-regulated in PCOS-like rats treated with metformin regardless of the occurrence of implantation. We also found that after metformin treatment PCOS-like rats with implantation exhibited decreased p-p38 MAPK, but not p38 MAPK, expression.

### Up-regulation of estrogen receptor (ER) expression in PCOS-like rats can be suppressed by metformin

Because estrogen-ER signaling regulates uterine PGR expression and activity (Li *et al.* 2014a; Patel *et al.* 2015) and because increased circulating E2 in PCOS-like rats can be inhibited by metformin treatment (Zhang *et al.* 2017), we sought to determine whether ER subtypes (ERα and ERβ) are involved in the regulation of aberrant PGR expression in PCOS-like rats and, if so, if metformin possibly alters ER subtype expression. Our data showed that PCOS-like rats exhibited increased *Esr1* (ERα) and *Esr2* (ERβ) mRNA levels, which were suppressed by metformin treatment. As shown in Figure 6A, while nuclear ERα immunoreactivity was detected in the epithelia and stroma in control rats treated with saline (Fig. 6B1), immunoreactivity of ERα was increased in the glandular epithelia and stroma in PCOS-like rats (Fig. 6D1). Furthermore, treatment with metformin led to decreased ERα immunoreactivity in control (Fig. 6C1) and PCOS-like rats (Fig. 6E1). No obvious difference in ERα immunoreactivity was observed in the myometrium in any of the groups (Fig. 6B2-E2). We also found that ERβ was mainly co-localized with ERα in the epithelia and stroma but not in the myometrium in control and PCOS-like rats regardless of the different treatments. Furthermore, with metformin treatment, we noted a significant increase in uterine *Esr1* and *Esr2* mRNAs in PCOS-like rats without implantation (Fig. 7A). Immunofluorescence staining revealed that, overall, immunoreactivities of both ERα and ERβ were diminished in the decidualizing stroma at the site of implantation (Fig. 7B1–D1), in the epithelia and stroma of the inter-implantation region (Fig. 7B2–D2) in control rats treated with saline or metformin (Fig. 7B2–C2), and in the inter-implantation site of PCOS-like rats treated with metformin (Fig. 7D2) compared to those rats before implantation (Fig 6. B1–E2). Interestingly, PCOS-like rats with no implantation exhibited sustained nuclear ERα immunoreactivity in the glandular epithelia and stroma (Fig. 7E1).

**Figure 6.**
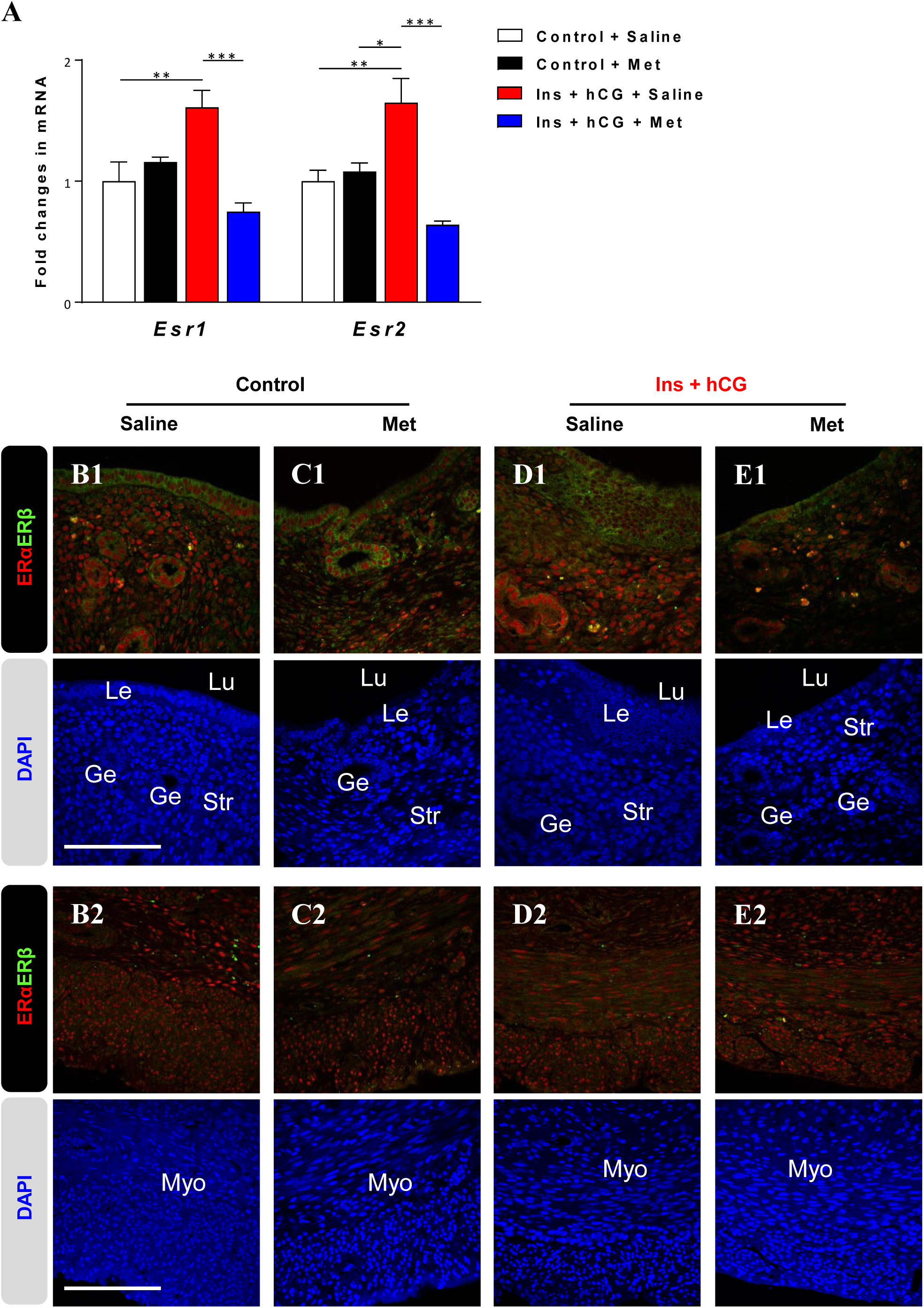
Chronic treatment with metformin alters ER subtype mRNA and protein expression in the rat uterus *in vivo*. A, Uterine tissues from control rats treated with saline vehicle or metformin and insulin+hCG-treated rats treated with saline or metformin (n = 6/group) were analyzed for mRNA levels of *Esr1* (ERα) and *Esr2* (ERβ) by qRT-PCR. The mRNA level of each gene relative to the mean of the sum of the *Gapdh* and *U87* mRNA levels in the same sample is shown. Values are expressed as means ± SEM. Statistical tests are described in the Material and Methods. * *p* < 0.05; ** *p* < 0.01; *** *p* < 0.001. B, Immunofluorescence detection of ERα (red) and ERβ (green) in control rats treated with saline (B1-2) or metformin (C1-2) and in insulin+hCG-treated rats treated with saline (D1-2) or metformin (E1-2). Representative images are shown (n = 5/group). Cell nuclei were counterstained with DAPI (blue, lower panel). Lu, lumen; Le, luminal epithelial cells; Ge, glandular epithelial cells; Str, stromal cells. Scale bars (100 µm) are indicated in the photomicrographs.

**Figure 7.**
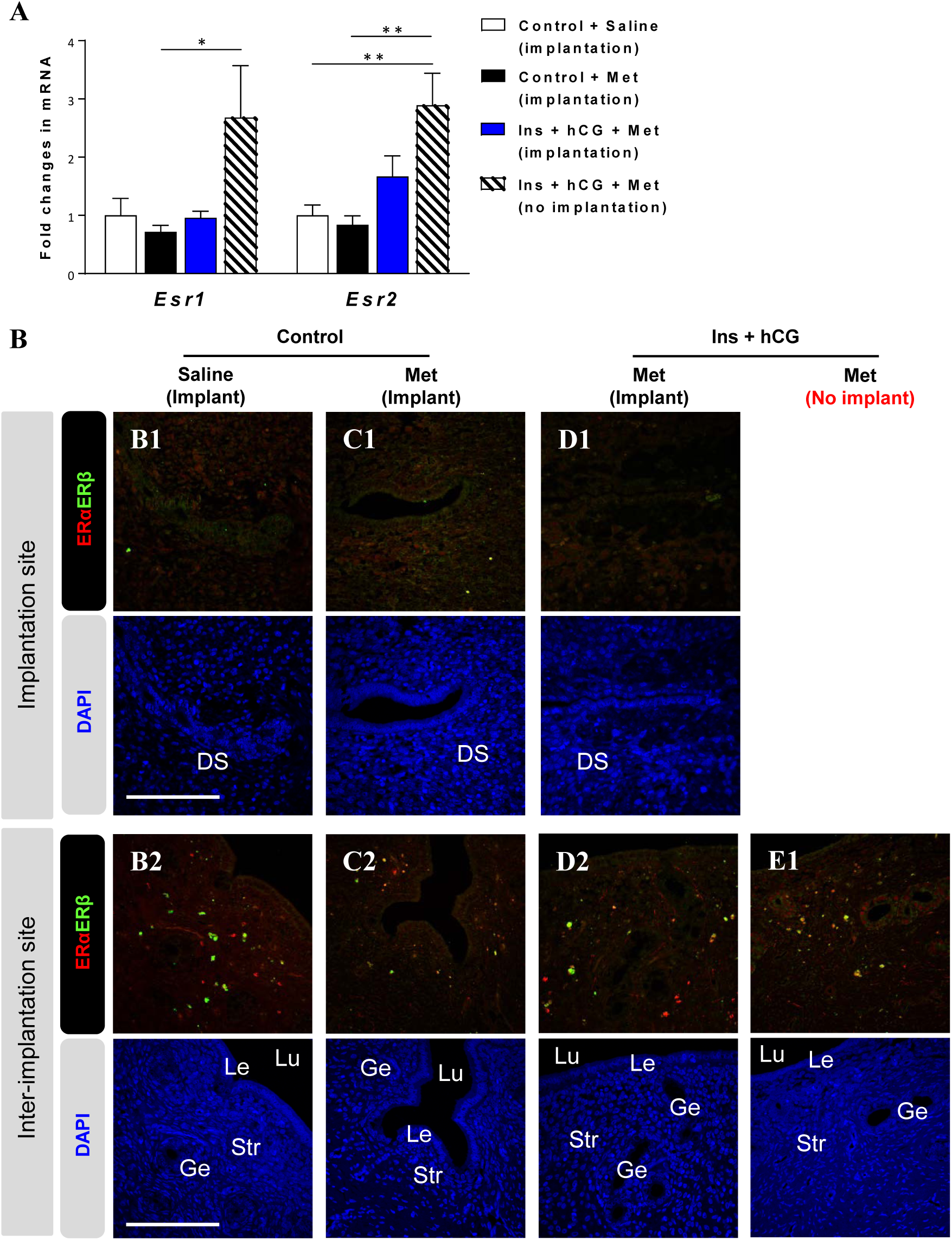
Chronic treatment with metformin alters ER subtype mRNA and protein expression in the rat uterus after implantation. A, Uterine tissues from control rats treated with saline or metformin and insulin+hCG-treated rats treated with saline or metformin (n = 5/group) were analyzed for mRNA levels of *Esr1* (ERα) and *Esr2* (ERβ) by qRT-PCR. The mRNA level of each gene relative to the mean of the sum of the *Gapdh* and *U87* mRNA levels in the same sample is shown. Values are expressed as means ± SEM. Statistical tests are described in the Materials and Methods. * *p* < 0.05; ** *p* < 0.01. B, Immunofluorescence detection of ERα (red) and ERβ (green) in control rats treated with saline (B1-2) or metformin (C1-2) and in insulin+hCG-treated rats treated with metformin with implantation (D1-2) or without implantation (E1). Representative images are shown (n = 5/group). Cell nuclei were counterstained with DAPI (blue, lower panel). DS, decidualized stroma; Lu, lumen; Le, luminal epithelial cells; Ge, glandular epithelial cells; Str, stromal cells. Scale bars (100 µm) are indicated in the photomicrographs.

### Differential cell-specific expression of phospho-histone H3 in PCOS-like rats treated with metformin

As previously demonstrated (Avellaira *et al.* 2006), p-histone H3 is of special interest because the endometrium of PCOS patients displays high levels of p-histone H3, which is associated with cellular processes such as mitosis (Brenner *et al.* 2003). Quantitative assessment of p-histone H3 indicated that no significant change in p-histone H3 immunoreactivity was present in the epithelia or stroma in any of groups (Supplemental Fig. 4E); however, metformin treatment decreased p-histone H3 immunoreactivity in the myometrium in PCOS-like rats compared to those treated with saline. Of note, intensely p-histone H3-positive stromal cells close to the luminal and glandular epithelia were found in PCOS-like rats treated with metformin (Supplemental Fig. 4D2). Similarly, p-histone H3 immunoreactivity was significantly increased in the stroma at the inter-implantation sites in PCOS-like rats treated with metformin independently of implantation (Supplemental Fig. 5E). In PCOS-like rats without implantation, p-histone H3 immunoreactivity was often detected in the luminal epithelia (Supplemental Fig. 5D1), although this was not statistically significant compared to PCOS-like rats with implantation (Supplemental Fig. 5E). It is thus likely that the regulation of mitotic activity by metformin is cell type-dependent in the uterus.

## Discussion

Reproductive dysfunction and infertility manifest noticeably in PCOS patients (Evans *et al.* 2016). In striking contrast to the attention given to hyperandrogenism and insulin resistance in women with PCOS, the aberrant P4 signaling pathway resulting in uterine P4 resistance has received much less attention (Li *et al.* 2014a; Piltonen 2016). This study is the first to show that the therapeutic effects of metformin on the regulation of uterine function in PCOS-like rats is mediated through P4 signaling.

Elucidating the regulation of endometrial PGR levels under PCOS conditions is important clinically. Our data show that increased PGR expression is paralleled with elevated ER expression in PCOS-like rats. This expression pattern is associated with an increased circulating E2 level (Zhang *et al.* 2016), suggests that E2-ER signaling contributes to the up-regulation of PGR under PCOS conditions *in vivo*. Similar to PCOS patients (Hu *et al.* submitted), PCOS-like rats also displayed high levels of PGR isoforms and ER subtypes in the uterus. The induction of implantation is required for the activation of PGR, and implantation subsequently alters gene expression in the endometrium (Gellersen and Brosens 2014; Patel *et al.* 2015); however, PGR-targeted gene expression in PCOS patients and PCOS-like rats has only been demonstrated to a limited degree. The current study shows that significantly decreased *Fkbp52* gene expression parallels increased expression of *Ihh, Smo*, and *Nr2f2* mRNAs without changes in *Ncoa2* mRNA in PCOS-like rats. In addition, we also found that abnormal expression of PGR-target genes, including *Fkbp52* and *Ncoa2*, is retained in PCOS-like rats with implantation failure. This is supported by *in vivo* studies showing that mice lacking *Fkbp52* (Tranguch *et al.* 2007; Yang *et al.* 2006) or *Ncoa2* (Mukherjee *et al.* 2007; Mukherjee *et al.* 2006) demonstrate the absence of decidualization after P4 supplementation due to diminished P4 responsiveness. Previous studies have reported that women with endometriosis and endometrial hyperplasia/carcinoma who develop P4 resistance have low levels of PGR expression (Gunderson *et al.* 2012; Shao *et al.* 2014a). Although it is currently unclear why differences exist in the regulation of uterine PGR expression between different diseases with P4 resistance, it is likely that uterine P4 resistance in PCOS-like rats is due to impaired PGR activity rather than PGR expression.

Defects in PGR isoform-specific P4 signaling in the mouse uterus can give rise to distinct phenotypes of uterine impairment and implantation failure (Li *et al.* 2014a). Here we observed no changes in total *Pgr* mRNA levels but a reduction of *Pgrb* mRNA levels in PCOS-like rat uterus after treatment with P4 and/or RU486. This suggests that uterine *Pgra* mRNA levels are increased. Meanwhile, several PGR target genes (e.g., *Ihh, Smo, Ptch, Nr2f2, Hand2*, and *Fkbp52*) are significantly altered in PCOS-like rats after P4 treatment. Thus, we speculate that changes in these P4-dependnent PGR target gene expression in PCOS-like rat uterus might be accounted for by an increase in *Pgra* mRNA expression. Studies of mutant mice lacking specific PGR isoform will clarify the functional differences between the two PGR isoforms in the progression of PCOS-induced uterine dysfunction.

P4-mediated and PGR-dependent regulation of ERK1/2 expression plays a critical role in humans and rodents during endometrial decidualization and implantation (Lee *et al.* 2013; Tapia-Pizarro *et al.* 2017; Thienel *et al.* 2002), but such regulation under PCOS conditions has not previously been reported. The inhibition of ERK1/2 expression and activation has been reported in ovarian granulosa and thecal cells in PCOS patients (Lan *et al.* 2015; Nelson-Degrave *et al.* 2005), and we have previously shown that the expression and activation of uterine ERK1/2 is suppressed in rats treated with insulin and hCG to induce the PCOS phenotype (Zhang *et al.* 2016). The present study supports and extends this work. Here we observed no changes in p-ERK1/2 or ERK1/2 expression in the rat uterus after prolonged treatment with insulin and hCG. However, we observed that the levels of p-c-Raf and p-MEK1/2, two upstream regulators of ERK1/2, were significantly decreased in PCOS-like rats after uterine implantation, establishing a tight link between different MAPK/ERK signaling molecules. Our data suggest that regulation of uterine ERK1/2 expression *in vivo* is time-dependent (Lee *et al.* 2013; Tapia-Pizarro *et al.* 2017; Thienel *et al.* 2002), which is similar to the regulation of PGR isoforms and PGR-targeted gene expression. The MAPK/ERK/p38 signaling pathway contributes to the regulation of inflammation and cytokine production (Arthur and Ley 2013; Cuadrado and Nebreda 2010), and the dysregulation of inflammation-related molecules is associated with PCOS conditions (Matteo *et al.* 2010; Orostica *et al.* 2016; Piltonen *et al.* 2013; Piltonen *et al.* 2015). Furthermore, like the activation of NFκB signaling that induces the transcriptional levels of inflammation-related gene expression in ovarian granulosa cells and in serum in PCOS patients (Liu *et al.* 2015; Zhao *et al.* 2015), our previous study has shown that the sustained metformin treatment markedly suppresses uterine inflammatory gene expression, especially the *Il-6* and *TNFα* mRNAs that are associated with inhibition of nuclear NF*к*B translocation in PCOS-like rats (Zhang *et al.* 2017). Importantly, p38 can antagonize ERK1/2 signaling mediated by protein phosphatase 2A and consequently down-regulate inflammatory cytokine and chemokine production (Cuadrado and Nebreda 2010), and the anti-inflammatory effects of MAPK/p38 are involved in the regulation of NF*к*B activity (Arthur and Ley 2013). These observations further indicate that metformin inhibits NF*к*B-driven inflammatory processes through p38 activation rather than through ERK1/2 inhibition in the PCOS-like rat uterus.

The results of the present study permit us to draw the following conclusions (Figure 8). 1) With sustained low levels of P4, the expressions of both uterine PGR isoforms are elevated in PCOS-like rats *in vivo*. This is positively associated with the high levels of ERs in PCOS-like rats. Consistent with mouse knockout studies, altered expression of *Fkbp52* and *Ncoa2*, two genes that contribute to uterine P4 resistance, is seen in PCOS-like rats before and after implantation. 2) Metformin directly suppresses uterine PGR isoform expression along with the correction of aberrant expression of PGR-targeted and implantation-related genes in PCOS-like rats. Abnormal cell-specific regulation of PGR and ER, paralleling the aberrant expression of PGR-targeted and implantation-related genes, is retained in PCOS-like rats with implantation failure. 3) Increased PGR expression is associated with inhibition of the MAPK/ERK/p38 signaling pathway, and the primary effect of metformin treatment is to restore the MAPK/p38 signaling pathway in the PCOS-like rat uterus. Taken together, our findings provide support for metformin therapy in the improvement of P4 signaling in PCOS-like rats with uterine dysfunction and for its clinical relevance in the treatment of PCOS patients with P4 resistance.

**Figure 8.**
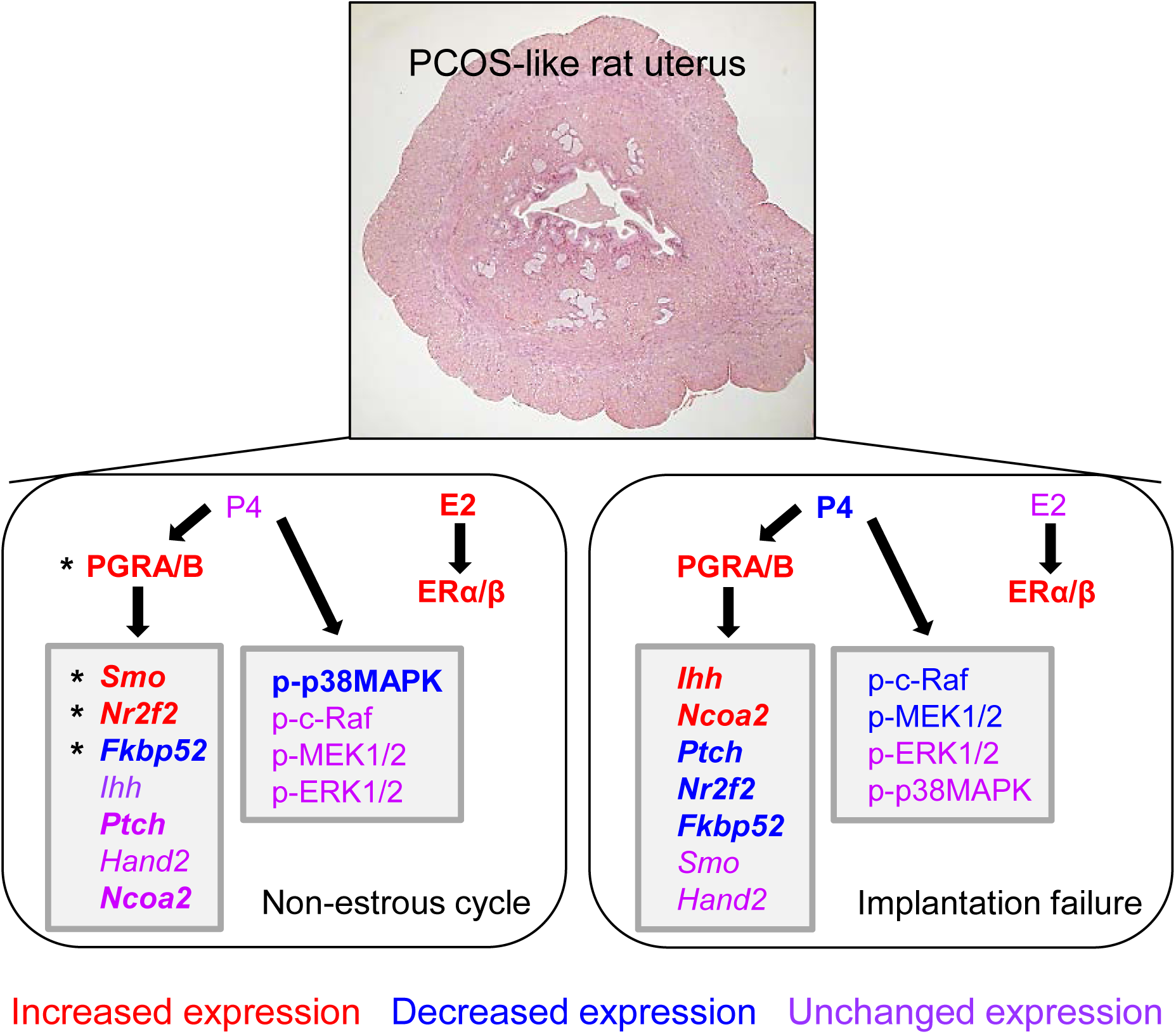
Schematic representation of the actions of metformin on the uterine progesterone signaling in the PCOS-like rats. Note that bold symbol indicates metformin-regulated genes and proteins and asterisk indicates that treatment with metformin or progesterone share the same targeted genes and proteins.

## Acknowledgements

The authors thank the Centre for Cellular Imaging at the University of Gothenburg and the National Microscopy Infrastructure (grant number VR-RFI 2016-00968) for providing assistance in microscopy.

## Materials and Methods

### Primary in vitro tissue culture and treatment

Uterine tissue culture and treatment was essentially carried out as described previously (Cui *et al.* 2015; Li *et al.* 2015). After rinsing in cold PBS, uterine tissues obtained from control (n = 5) and PCOS-like (n = 5) rats were divided into equal-sized explants and placed in 24-well tissue culture plates (Sarstedt, Newton, MA) containing RPMI-1640 medium with charcoal-stripped 10% fetal calf serum and 100 IU/ml penicillin/streptomycin (GIBCO-BRL, San Francisco, CA). Cultured tissues treated with sterile saline or metformin (10 mM in sterile saline) were incubated in a humidified incubator (37°C, 95% O_2_, 5% CO_2_) and separately collected at 0 h, 24 h, 48 h, and 72 h after treatment (Supplemental Fig. 2). Each culture condition was performed in five replicates (five wells), and tissues from a minimum of five control rats and five PCOS-like (insulin+hCG-treated) rats were used. At the end of the experiments, cultured tissues were snap-frozen in liquid nitrogen and stored at –70°C.

### Morphological assessment and immunostaining

Uterine tissues were fixed in 10% formalin, embedded in paraffin, and sectioned for hematoxylin and eosin staining according to standard procedures. Immunohistochemistry and immunofluorescence were performed according to previously described methods (Zhang *et al.* 2017; Zhang *et al.* 2016). The endogenous peroxidase and nonspecific binding were removed by incubation with 3% H_2_O_2_ for 10 min and with 10% normal goat serum for 1 h at room temperature. After incubation with the primary antibody (Supplemental Table 1) overnight at 4°C in a humidified chamber, the sections were stained using the avidin-biotinylated-peroxidase complex detection system (Vector Laboratories Inc., Burlingame, CA) followed by treatment with 3-amino-9-ethyl carbazole developing reagent plus High Sensitivity Substrate (SK-4200, Vector Laboratories). The sections were imaged on a Nikon E-1000 microscope (Japan) and photomicrographed using Easy Image 1 (Bergström Instrument AB, Sweden).

The other half of the uterine sections were incubated with primary antibody in 0.01 M Tris-buffered saline supplemented with Triton X-100 (TBST) containing 5% nonfat milk overnight at 4°C, and a secondary antibody was applied at room temperature for 1 h. After the sections were washed with TBST, they were re-suspended in mounting medium containing DAPI (4′,6′-diamidino-2-phenylindole; Vector Laboratories). Sections were examined under an Axiovert 200 confocal microscope (Zeiss, Jena, Germany) equipped with a laser-scanning confocal imaging LSM 710 META system (Carl Zeiss) and photomicrographed. Background settings were adjusted from the examination of negative control specimens. Images of positive staining were adjusted to make optimal use of the dynamic range of detection. The immune staining was quantified by semi-quantitative histogram scoring (Q-H score) as described previously (Mariee *et al.* 2012). All morphological and immunohistochemical assays were performed by at least two researchers in an operator-blinded manner.

### Protein isolation and Western blot analysis

A detailed explanation of the Western blot analysis protocol has been published (Zhang *et al.* 2017; Zhang *et al.* 2016). Total protein was isolated from whole uterine tissue by homogenization in RIPA buffer (Sigma-Aldrich) supplemented with cOmplete Mini protease inhibitor cocktail tablets (Roche Diagnostics, Mannheim, Germany) and PhosSTOP phosphatase inhibitor cocktail tablets (Roche Diagnostics). After determining total protein by Bradford protein assay, equal amounts (30 µg) of protein were resolved on 4–20% TGX stain-free gels (Bio-Rad Laboratories GmbH, Munich, Germany) and transferred onto PVDF membranes. The membranes were probed with the primary antibody (Supplemental Table 1) in TBST containing 5% non-fat dry milk followed by HRP-conjugated secondary antibody. When necessary, the PVDF membranes were stripped using Restore PLUS Western blot stripping buffer (Thermo Scientific, Rockford, IL) for 15 minutes at room temperature, washed twice in TBST, and then re-probed. Ultraviolet activation of the Criterion stain-free gel on a ChemiDoc MP Imaging System (Bio-Rad) was used to control for proper loading. Band densitometry was performed using Image Laboratory (Version 5.0, Bio-Rad).

In the present study, we used a novel stain-free technology – ultraviolet activation of the Criterion stain-free gel on a ChemiDoc MP Imaging System (Bio-Rad) – which was used to control proper loading (total protein normalization). This technology represents a significant advancement over existing stain-based total protein quantitation approaches such as Coomassie blue, Ponceau S, and others and gives accurate protein loading control data in a standardized manner without requiring lengthy optimization. The detection of proteins on stain-free gels is based upon trihalocompound modification of tryptophan residues in proteins run on stain-free gels, which are exposed to UV. The modified tryptophans give a fluorescent signal that can be readily detected by a CCD camera. In our study, band densitometry was performed using Image Laboratory. When quantified, the intensity of each protein band was normalized to the total protein in individual sample. This method has been commonly used in both human and animal tissues to semi-quantify concentrations of specific proteins in many studies. Please visit the link given below for review of previous publications using the same method and technology:

(http://www.biorad.com/webroot/web/pdf/lsr/literature/Bulletin_6351.pdf).

### RNA extraction and qRT-PCR analysis

For RNA isolation, tissues from each rat were lysed using TRIzol Reagent (Life Technologies), and RNA was isolated following standard protocols. qRT-PCR was performed with a Roche Light Cycler 480 sequence detection system (Roche Diagnostics Ltd., Rotkreuz, Switzerland) as previously described (Zhang *et al.* 2017; Zhang *et al.* 2016). The PCR amplifications were performed with a SYBR green qPCR master mix (#K0252, Thermo Scientific, Rockford, IL). Total RNA was prepared from the frozen whole uterine tissues, and single-stranded cDNA was synthesized from each sample (2 µg) with M-MLV reverse transcriptase (#0000113467, Promega Corporation, Fitchburg, WI) and RNase inhibitor (40 U) (#00314959, Thermo Scientific). cDNA (1 µl) was added to a reaction master mix (10 µl) containing 2× SYBR green qPCR reaction mix (Thermo Scientific) and gene-specific primers (5 µM each of forward and reverse primers). All primers are indicated in Supplemental Table 2. All reactions were performed six times, and each reaction included a non-template control. The results for target genes were expressed as the amount relative to the average CT values of *GAPDH* + *U87* in each sample. Relative gene expression was determined with the 2–ΔΔCT method, and the efficiency of each reaction – as determined by linear regression – was incorporated into the equation.

### Measurement of biochemical parameters

Concentrations of gonadotropins (follicle stimulating hormone and luteinizing hormone), steroid hormones (17β-estradiol, progesterone, testosterone, 5α-dihydrotestosterone, and androstenedione), sex hormone-binding globulin, glucose, and insulin in rat serum samples were measured using commercially available assays (Cloud-Clone Corp., Houston, TX) as described previously (Zhang et al. 2017). All samples and standards were measured in duplicate. The intra- and inter-assay coefficients of variation are listed in Supplemental Table 3.

---

**Supplementary Table 1.** Antibodies: species, clone/catalog number, method, dilution, and source.

| Antibody | Species | Cat. No. | kDa | Method | Dilution | Source |
| --- | --- | --- | --- | --- | --- | --- |
| PGR | Mouse | MA1-410 | B120 | WB | 1:500 | Thermo Fisher (Rockford, IL) |
|  |  |  | A94 | IHC | 1:50 |  |
| p-c-Raf | Rabbit | 9427 | 74 | WB | 1:500 | Cell Signaling Technology<br>(Danver, MA) |
| p-MEK1/2 | Rabbit | 9154 | 45 | WB | 1:500 | Cell Signaling Technology |
| p-ERK1/2 | Rabbit | 4370 | 42,44 | WB | 1:1000 | Cell Signaling Technology |
| ERK1/2 | Rabbit | 4695 | 42,44 | WB | 1:2000 | Cell Signaling Technology |
| p-p38 MAPK | Rabbit | 4511 | 43 | WB | 1:300 | Cell Signaling Technology |
| p38 MAPK | Rabbit | 8690 | 40 | WB | 1:1000 | Cell Signaling Technology |
| p-JNK | Rabbit | 4668 | 46,54 | WB | 1:500 | Cell Signaling Technology |
| JNK | Rabbit | 9252 | 46,54 | WB | 1:1000 | Cell Signaling Technology |
| ERα | Mouse | 6F11 | | IHC, IF | 1:50 | Novocastra Laboratories Ltd.<br>(Newcastle upon Tyne, UK) |
| ERβ | Rabbit | Ab3577 | | IF | 1:50 | Abcam (Cambridge, UK) |
| p-histone H3<br>(Ser10) | Rabbit | 3377 |  | IF | 1:100 | Cell Signaling Technology |

PGR, progesterone receptor; p-MEK1/2, phosphorylation-MAP2K1/2; ERK1/2, extracellular signal-regulated kinases 1/2; p38 MAPK, p38 mitogen-activated protein kinase; JNK, jun-amino-terminal kinase; ERα , estrogen receptor alpha; WB, western blot; IHC, immunohistochemistry; IF, immunofluorescence.

**Supplementary Table 2.**
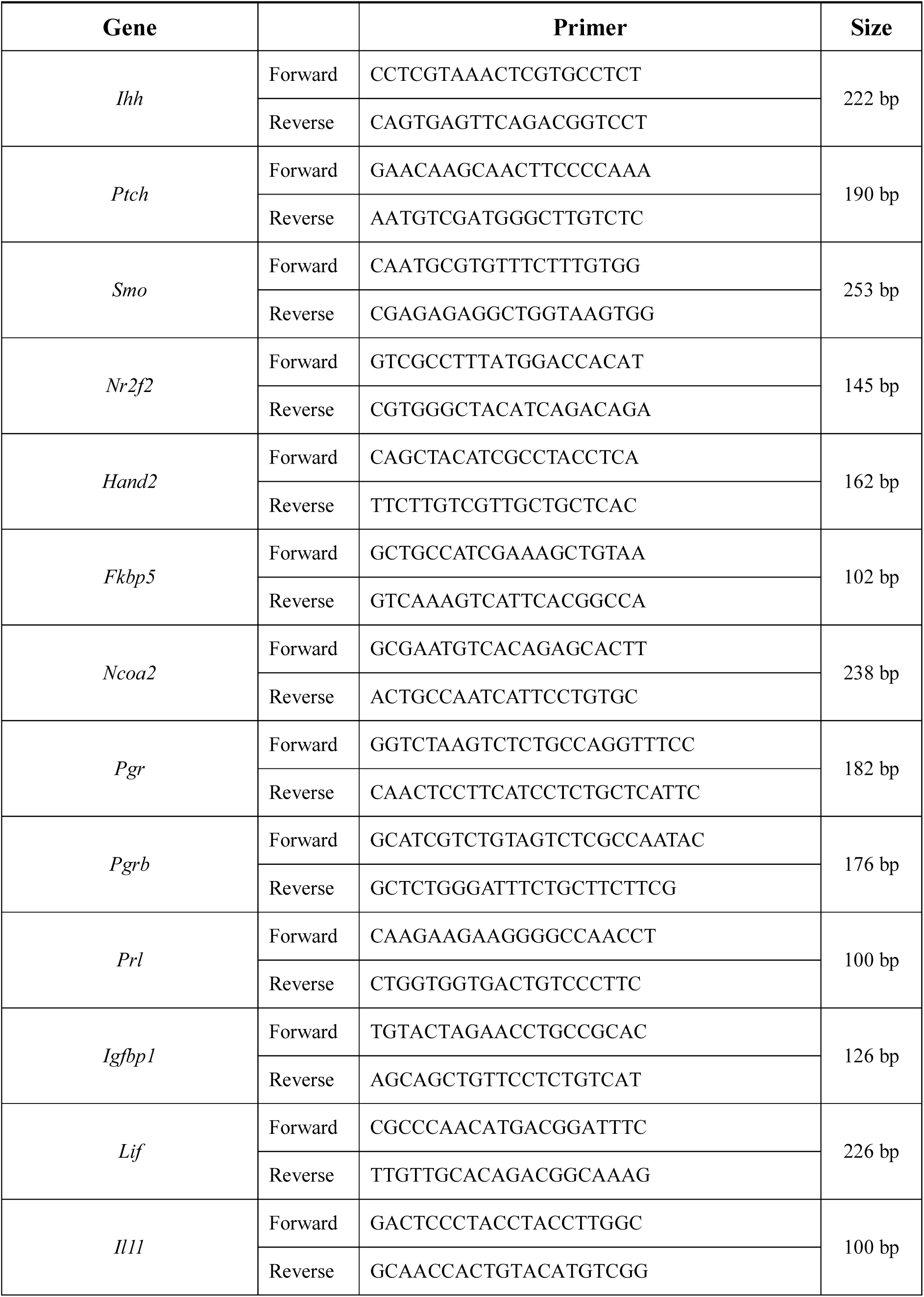
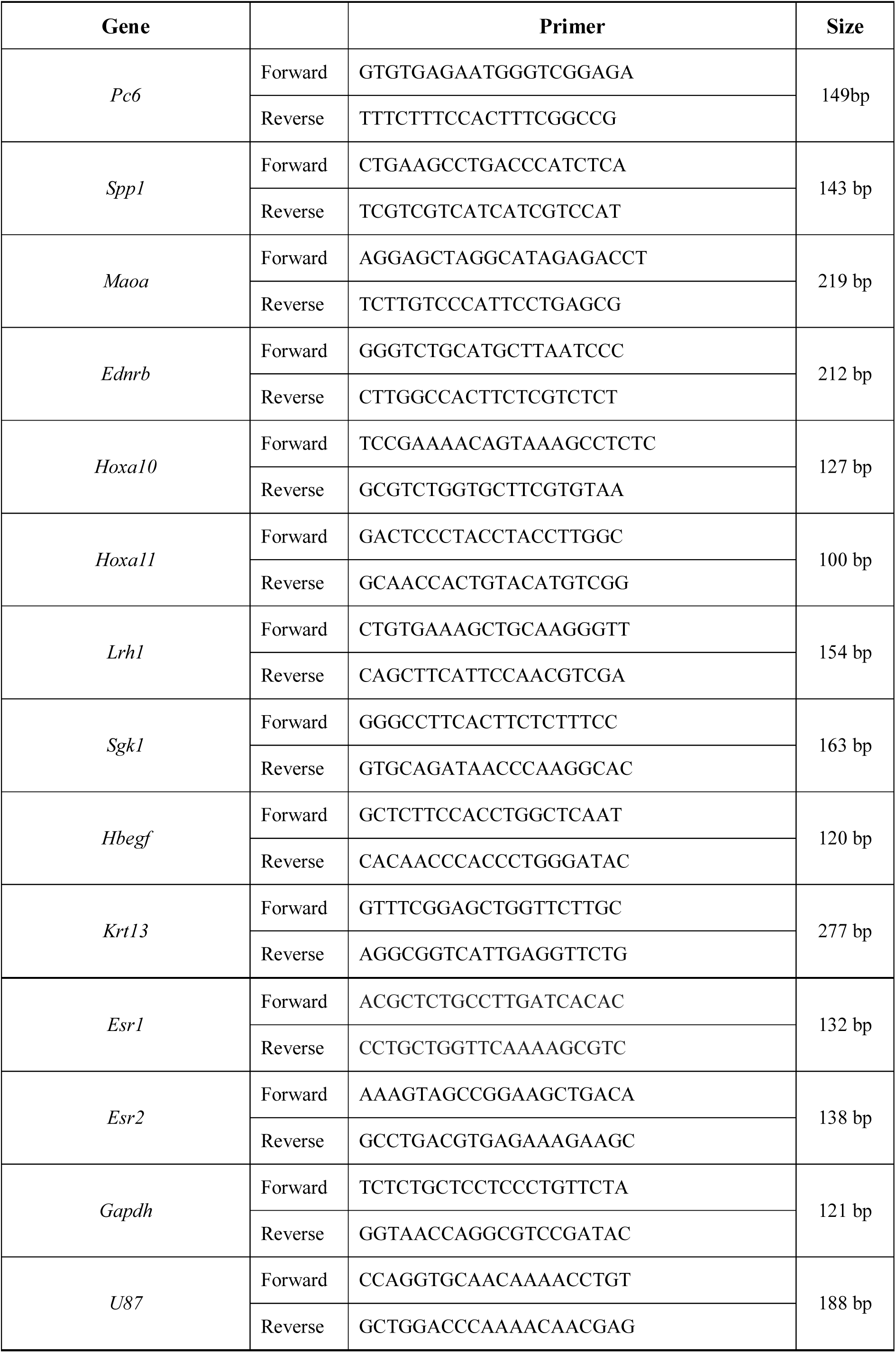
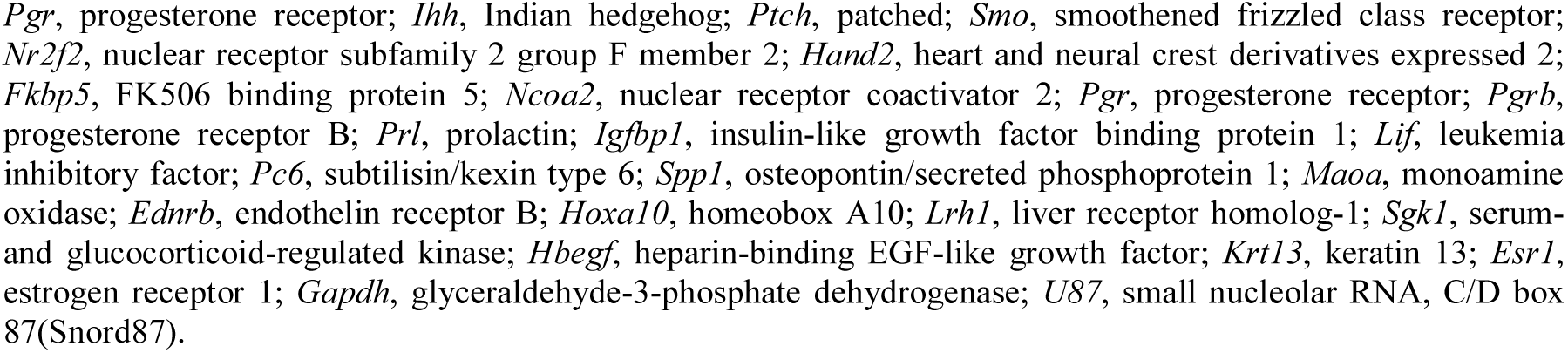
Sequences of primer pairs used for qRT-PCR measurement.

**Supplementary Table 3.** The intra-assay and inter-assay % CV for the rat gonadotropins, steroid hormones, and insulin.

|  | Intra-assay % CV | Inter-assay % CV |
| --- | --- | --- |
| FSH | 6.4 | 6.8 |
| LH | 6.4 | 6.6 |
| E2 | 6.5 | 6.8 |
| P4 | 6.4 | 6.6 |
| Total T | 6.2 | 6.6 |
| A4 | 6.7 | 6.9 |
| DHT | 6.2 | 6.7 |
| SHBG | 6.4 | 6.8 |
| Insulin | 6.3 | 6.6 |

FSH, follicle-stimulating hormone; LH, luteinizing hormone; E2, 17β-estradiol; P4, progesterone; T, testosterone; A4, androstenedione; DHT, 5α-dihydrotestosterone; SHBG, sex hormone-binding globulin.

**Supplementary Table 4.** Effects of implantation on the endocrine and metabolic alterations in control rats treated with saline or metformin, and insulin+hCG-treated (PCOS-like) rats with metformin.

|  | **Saline**<br>(n = 8) | **Met**<br>(n = 10) | **Insulin + hCG**<br>**+ Met**<br>**(implantation)**<br>(n = 10) | **Insulin + hCG**<br>**+ Met**<br>**(no implantation)**<br>(n = 5) |
| --- | --- | --- | --- | --- |
| FSH (ng/ml) | 2.13 ± 0.04 | 2.06 ± 0.04 | 2.14 ± 0.04 | 2.14 ± 0.10 |
| LH (ng/ml) | 2.00 ± 0.24 | 1.93 ± 0.11 | 2.31 ± 0.17 | 2.73 ± 0.36 |
| LH / FSH | 0.94 ± 0.11 | 0.94 ± 0.05 | 1.08 ± 0.08 | 1.28 ± 0.18 |
| E2 (ng/ml) | 1.31 ± 0.04 | 1.34 ± 0.03 | 1.23 ± 0.03 | 1.32 ± 0.05 |
| P4 (ng/ml) | 6.88 ± 0.35 | 6.51 ± 0.37 | 8.95 ± 0.44 <sup>a, c</sup> | 4.80 ± 0.11 <sup>b, d, e</sup> |
| Total T (ng/ml) | 5.11 ± 0.71 | 3.83 ± 0.22 | 3.78 ± 0.44 | 7.40 ± 0.09 <sup>a, c, e</sup> |
| A4 (ng/ml) | 0.142 ± 0.004 | 0.142 ± 0.006 | 0.162 ± 0.006 | 0.181 ± 0.016 <sup>b, c</sup> |
| DHT (pg/ml) | 102.97 ± 5.66 | 97.92 ± 9.28 | 102.00 ± 5.52 | 154.58 ± 8.39 <sup>a, c, e</sup> |
| E2 / Total T | 0.29 ± 0.04 | 0.36 ± 0.02 | 0.38 ± 0.05 | 0.18 ± 0.01 <sup>d, f</sup> |
| Total T / DHT | 0.051 ± 0.007 | 0.044 ± 0.006 | 0.038 ± 0.005 | 0.049 ± 0.003 |
| Total T / A4 | 36.37 ± 5.36 | 27.28 ± 1.77 | 23.26 ± 2.49 | 42.20 ± 3.52 <sup>d, e</sup> |
| SHBG (ng/ml) | 11.08 ± 0.33 | 10.89 ± 0.35 | 9.67 ± 1.16 | 10.91 ± 0.56 |
| Fasting glucose (mmol/l) | 4.19 ± 0.09 | 3.92 ± 0.10 | 4.53 ± 0.19 <sup>d</sup> | 4.98 ± 0.12 <sup>a, c</sup> |
| Fasting insulin (pg/ml) | 3.26 ± 0.05 | 3.17 ± 0.02 | 3.22 ± 0.04 | 3.42 ± 0.03 <sup>b, c, f</sup> |
| HOMA-IR | 0.61 ± 0.01 | 0.55 ± 0.01 | 0.65 ± 0.03 <sup>c</sup> | 0.76 ± 0.02 <sup>a, c, f</sup> |

Met, metformin; FSH, follicle-stimulating hormone; LH, luteinizing hormone; E2, 17 $\alpha$ -estradiol; P4, progesterone; T, testosterone; A4, androstenedione; DHT, 5α-dihydrotestosterone; SHBG, sex hormone-binding globulin; HOMA-IR, Homeostasis model assessment of insulin resistance, HOMA-IR= fasting blood glucose (mmol/l) × fasting serum insulin ( $\mu$ U/ml) / 22.5.
Values are Mean ± SEM. The multiple comparisons between data were performed using one-way ANOVA and Tukey's post hoc test. A P-value less than 0.05 was considered statistically significant.
<sup>a</sup> P < 0.01 versus Saline group.
<sup>b</sup> P < 0.05 versus Saline group.
<sup>c</sup> P < 0.01 versus Met group.
<sup>d</sup> P < 0.05 versus Met group.
<sup>e</sup> P < 0.01 versus Insulin + hCG + Met (implantation) group.
<sup>f</sup> P < 0.05 versus Insulin + hCG + Met (implantation) group.

**Supplementary Figure 1.**
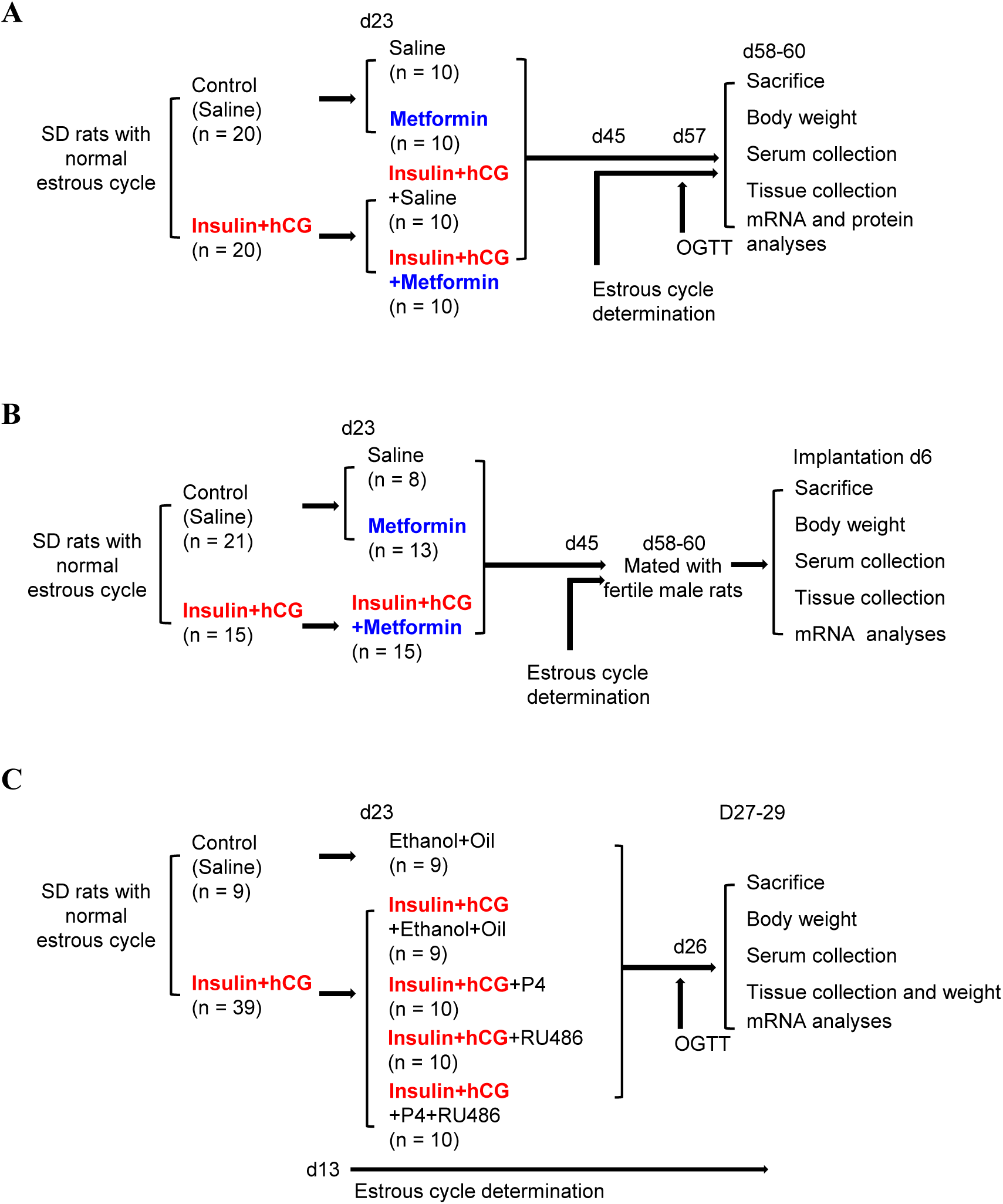
Schematic representation of experimental groups and treatment protocol. In, the final number of animals.

**Supplementary Figure 2.**
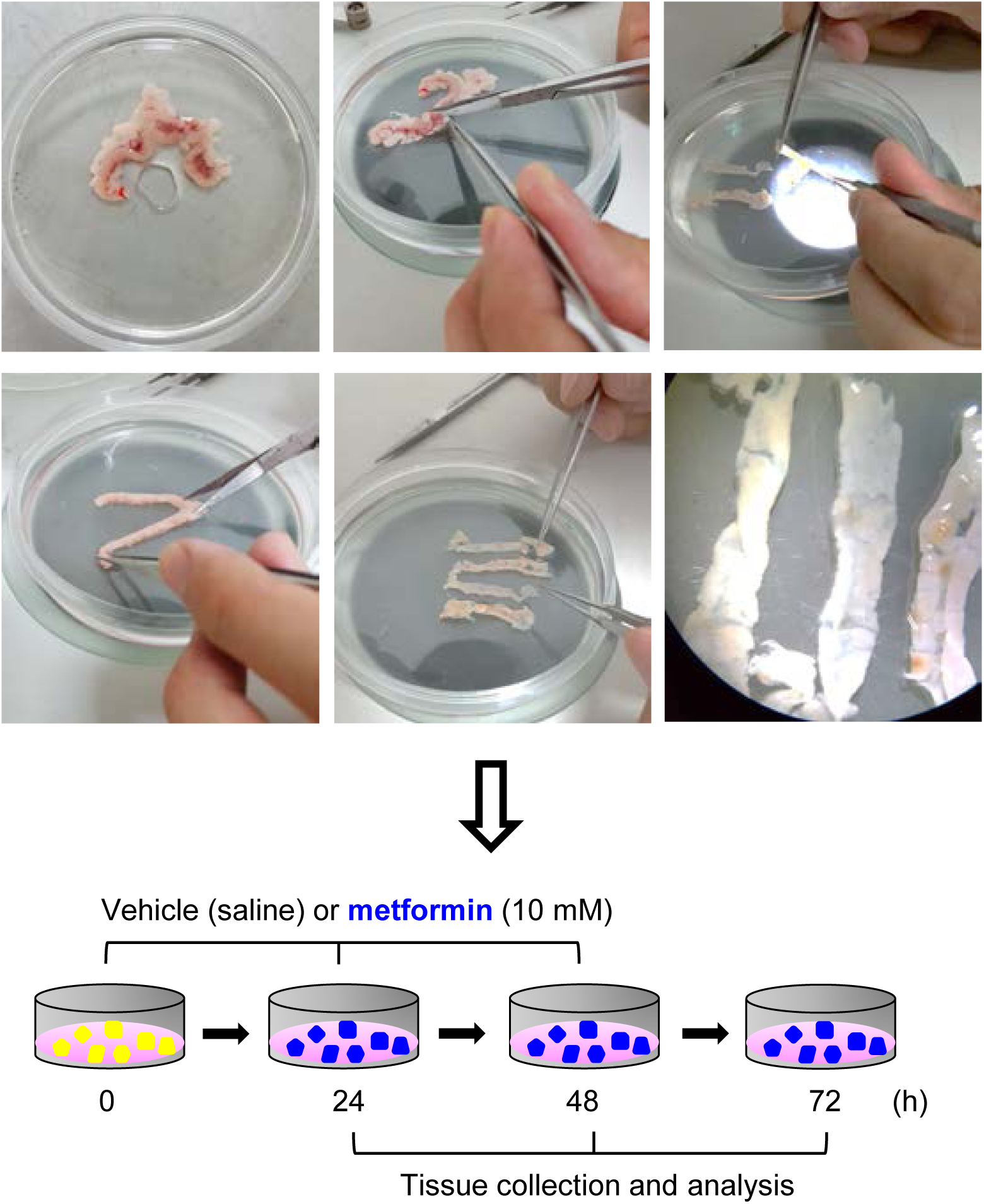
Photographs and illustration of the procedures for rat uterine tissue preparation and culture with and without metformin. Culture media with and without metformin treatment were changed daily.

**Supplementary Figure 3.**
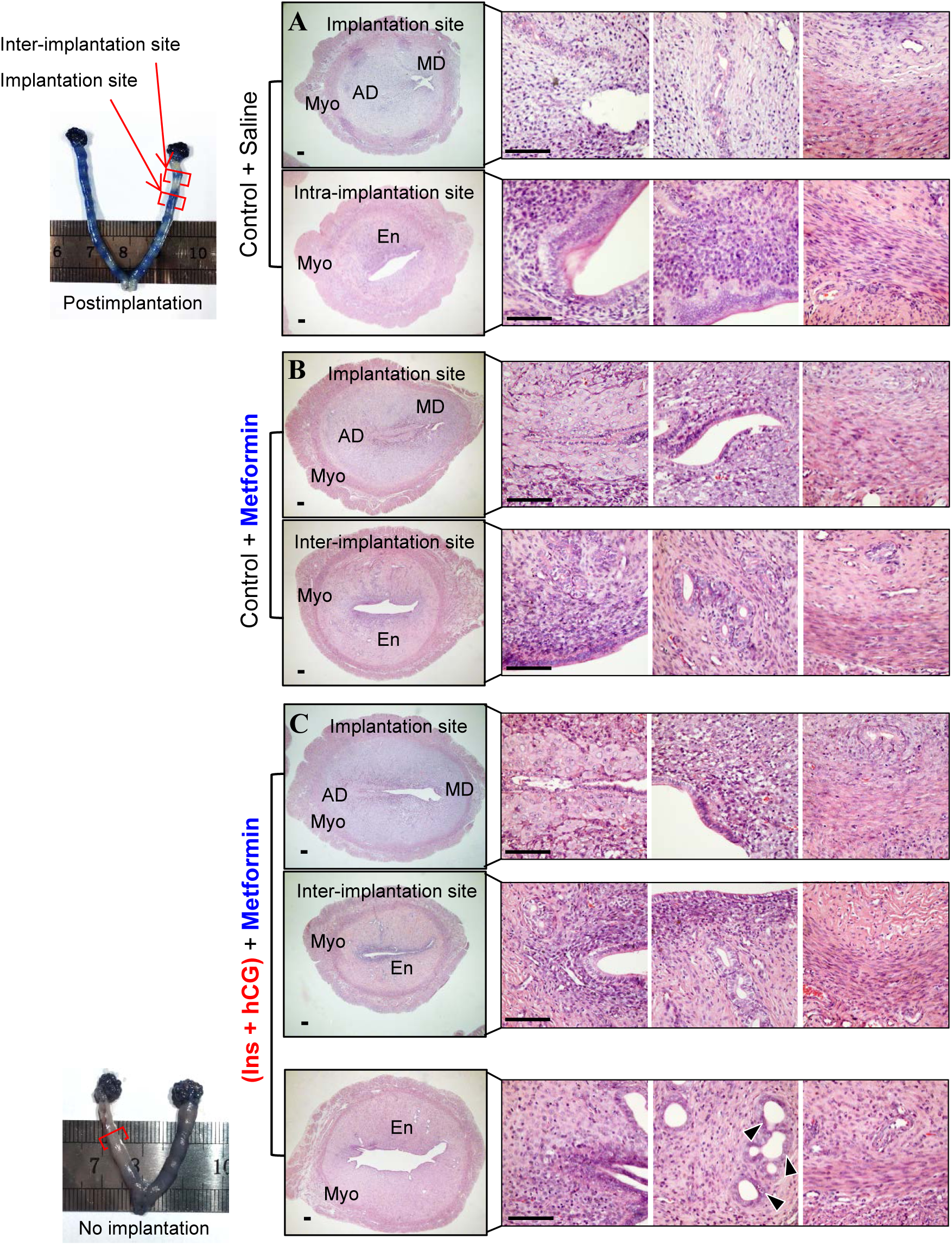
Effects of metformin on uterine structure in rats after implantation. Female rats were mated individually with fertile male rats according to their estrous cycle stage. Representative photomicrographs of uteri collected on day 6 after pregnancy with implantation sites visualized by Chicago Blue B dye injection. Because the insulin+hCG-treated rats without a normal estrous cycle display implantation failure, these rats treated with saline were excluded from the analysis. Higher-magnification images of the different areas are shown on the rightmost three panels of each row. MD, mesometrial decidua; AD, antimesometrial decidua; En, endometrium; Myo, myometrium. Scale bars (100µm) are indicated in the photomicrographs.

**Supplementary Figure 4.**
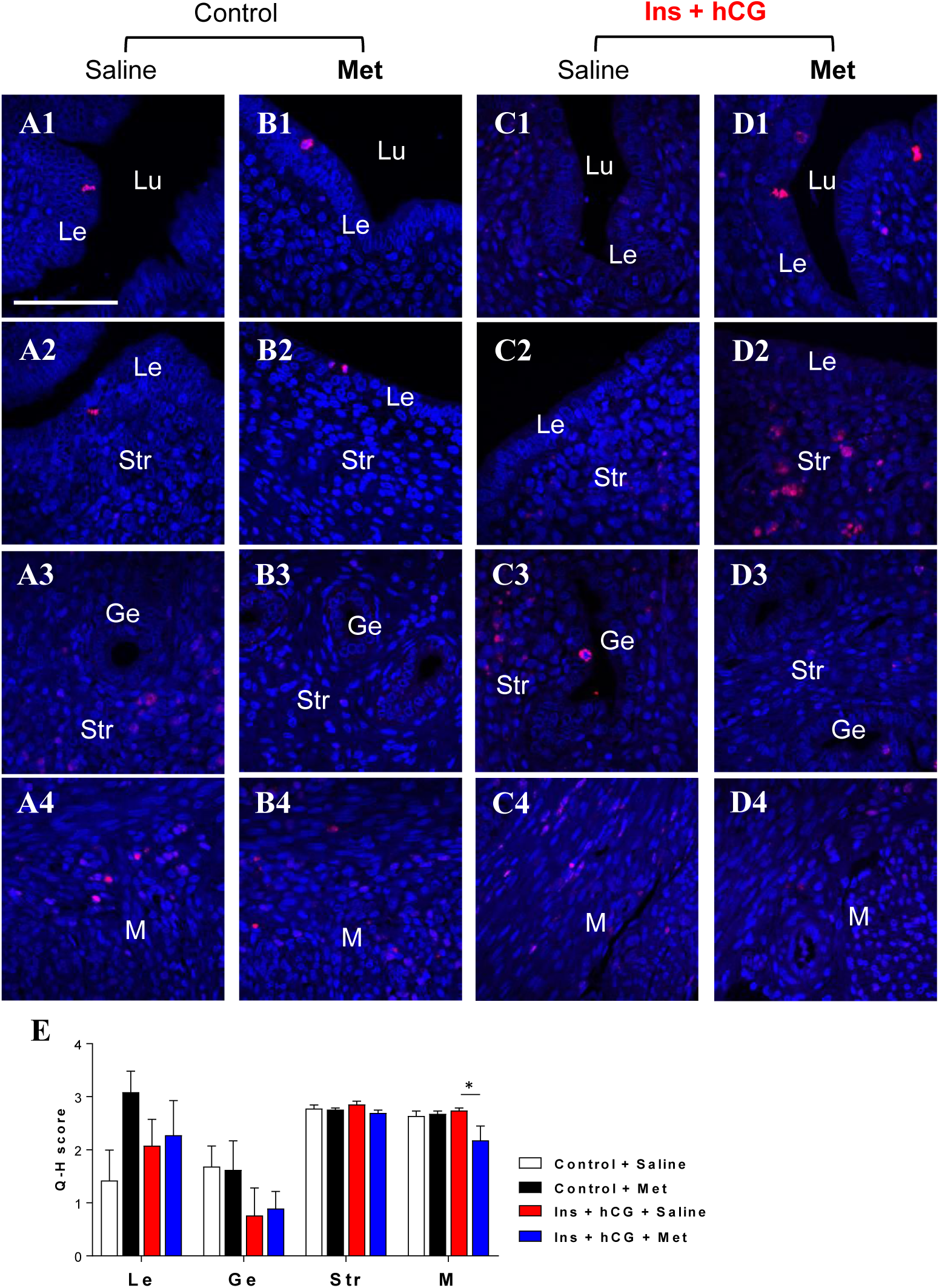
Chronic treatment with metformin alters p-histone H3 (Ser10) protein expression in the rat uterus *in vivo*. Immunofluorescence detection of p-histone H3 (red) in control rats treated with saline (A1-4) or metformin (B1-4) and in insulin+hCG-treated rats treated with saline (C1-4) or metformin (D1-4). Cell nuclei were counterstained with DAPI (blue). Lu, lumen; Le, luminal epithelial cells; Ge, glandular epithelial cells; Str, stromal cells; M, myometrium. Scale bars (100 µm) are indicated in the photomicrographs. The number of all p-histone H3-positive cells in whole luminal epithelial cells, glandular epithelial/stromal cells and myometrium)/uterine section/animal was counted. Semi-quantitative immunofluorescence of p-histone H3 (Q-H score) in the different uterine cell types (n = 5/group) is shown in E. Values are expressed as means ± SEM. Statistical tests are described in the Material and Methods. * *p* < 0.05.

**Supplementary Figure 5.**
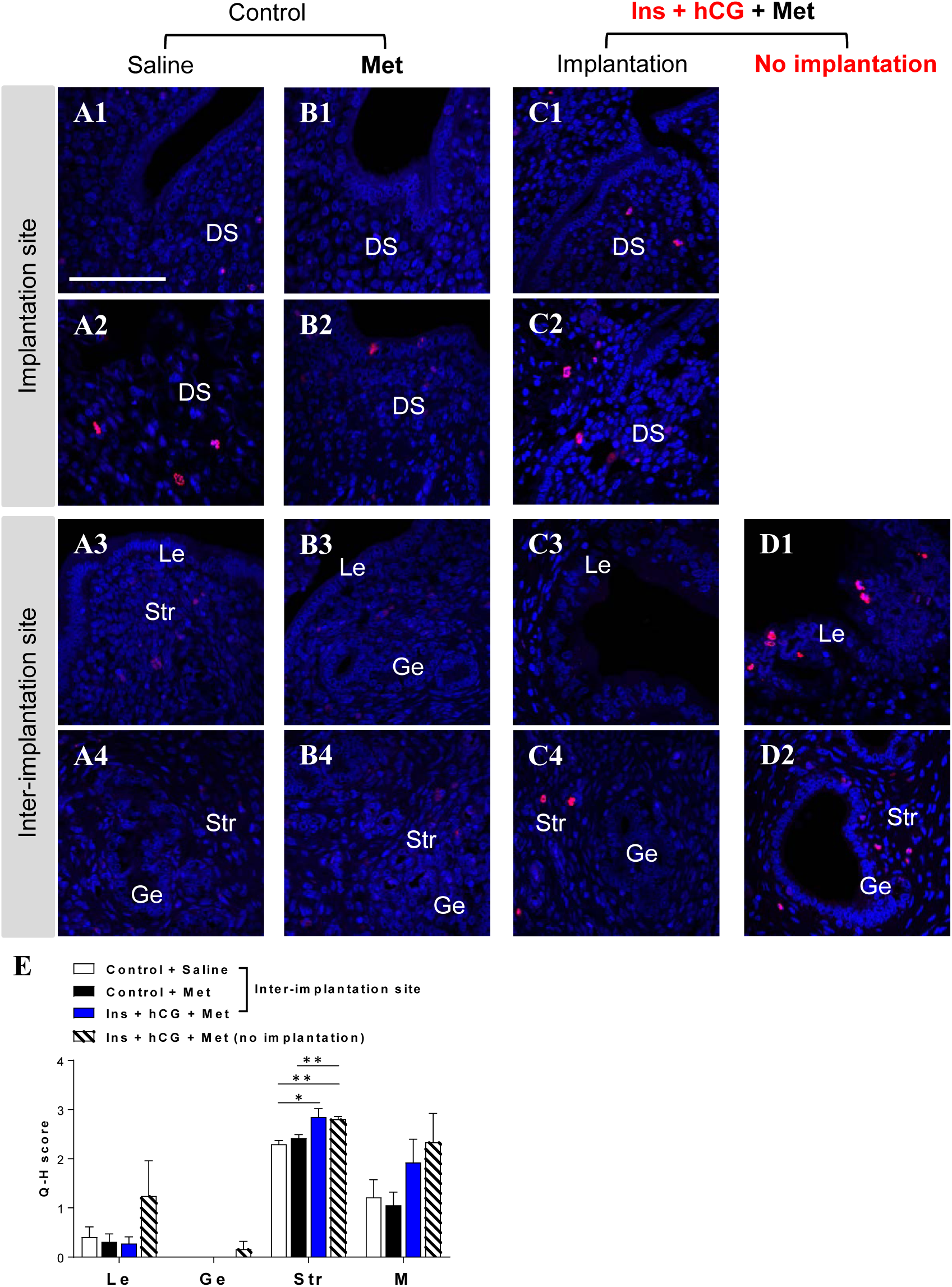
Chronic treatment with metformin alters p-histone H3 (Ser10) protein expression in the rat uterus after implantation. Immunofluorescence detection of p-histone H3 (red) in control rats treated with saline (A1-4) or metformin (B1-4) and in insulin+hCG-treated rats treated with metformin with implantation (C1-4) or without implantation (D1-2). Cell nuclei were counterstained with DAPI (blue). DS, decidualized stroma; Le, luminal epithelial cells; Ge, glandular epithelial cells; Str, stromal cells; M, myometrium. Scale bars (100 µm) are indicated in the photomicrographs. The number of all p-histone H3-positive cells in whole luminal epithelial cells, glandular epithelial/stromal cells and myometrium)/uterine section/animal was counted. Semi-quantitative immunofluorescence of p-histone H3 (Q-H score) in the different uterine cell types is shown in E. Uterine tissues were analyzed (n = 8/group; n = 5 for insulin+hCG-treated rats with metformin but without implantation). Values are expressed as means ± SEM. Statistical tests are described in the Materials and Methods. * *p* < 0.05; ** *p* < 0.01.

